# The human gut virome is a non-redundant and clinically informative component of the microbiome

**DOI:** 10.64898/2026.05.13.724676

**Authors:** Yiyan Yang, Dan Huang, Joshua R. Korzenik, Scott T. Weiss, Yang-Yu Liu, Zheng Sun

**Author notes:** Correspondence: Y.-Y.L.; Z.S.

## Abstract

The gut virome represents a vast reservoir of genetic diversity with profound implications for human health, yet it remains the “dark matter” of the microbiome due to the staggering complexity of reproducible viral profiling. It remains fundamentally contested whether biologically informative virome signals can be robustly recovered from routine whole-metagenome sequencing (WMS), and to what extent these signals offer ecological insights independent of the bacteriome. Here we present VIP2B, a framework that leverages Type IIB restriction tags to extract multifaceted viral features (encompassing taxonomy, coverage, function, and phenotype) directly from bulk WMS data. Through extensive benchmarking across incomplete references, unseen genomes, and high bacterial or host background, we demonstrate that VIP2B achieved high precision and robust taxonomic concordance. By applying VIP2B to paired bulk and virus-like particle (VLP)-enriched datasets, we reveal a species-level overlap far greater than previously recognized, proving that standard bulk metagenomes contain a wealth of recoverable viral information. Analysis of 20 clinical cohorts demonstrates that coverage-, function-, and phenotype-resolved viral features consistently identify disease-associated signatures that escape taxonomic analysis alone, significantly improving diagnostic models over bacteriome-only approaches. Finally, we define two distinct gut virome community states at the population scale (n=6,090), characterized by divergent diversity profiles and health associations. Our findings establish the gut virome as a non-redundant, clinically actionable component of the human holobiont and provide the methodology necessary to transition microbiome research toward a truly multi-kingdom framework.

## INTRODUCTION

Viruses are the most abundant biological entities in the biosphere, and wherever bacteria thrive, viruses that infect bacteria (bacteriophages or phages) impose a primary layer of ecological control^1–4^. Through density-dependent predation, lysogenic integration, and horizontal gene transfer, phages can rapidly restructure bacterial populations and reprogram bacterial traits, with downstream effects on community metabolism, colonization resistance, and the dissemination of virulence- or fitness-associated genes^5–8^. The human gut is no exception: its virome is typically phage-dominated and tightly interwoven with dense bacterial consortia, and shifts in virome composition and functional potential have been linked to disease-associated ecosystem remodeling, including inflammatory bowel disease^9–13^ and metabolic syndrome^14–17^, yet several fundamental questions remain unresolved. How much of the gut virome can be reliably recovered from routine whole-metagenome sequencing (WMS) rather than dedicated virus-like particle (VLP)-enriched sequencing^18^? Does the virome provide biologically meaningful information beyond what is already captured by the bacteriome? Are disease-associated virome perturbations adequately described by taxonomic abundance alone, or do they instead extend across broader functional, ecological, and community-level dimensions? And, at the population scale, does the gut virome exhibit recurrent organizational states linked to health and disease? Addressing these questions is essential for determining whether the virome should be treated as an independent component of microbiome analysis rather than as a technically challenging adjunct. Yet, progress on these questions has been limited by the lack of scalable, standardized approaches to virome profiling.

In contrast to bacteriome profiling, virome profiling is inherently more challenging because viruses lack broadly conserved universal marker genes, making simple, lineage-independent counting (“DNA-to-Marker strategy”) difficult across diverse lineages^19,20^. Consequently, many gut virome studies have relied on assembly-first workflows^21,22^: assemble contigs, curate viral sequences, and quantify by read mapping. Assembly remains indispensable for discovering previously uncharacterized viruses, but it is computationally intensive, sensitive to sequencing depth and parameter choices, and can under-represent low-abundance genomes, all of which complicate standardization across datasets. These limitations have renewed interest in assembly-free, reference-based profiling^23,24^, where reads are assigned directly to whole-genome viral catalogs (“DNA-to-DNA strategy”). This approach is now increasingly practical because gut viral reference resources have expanded rapidly, including recently published, systematically organized catalogs such as the Unified Human Gut Virome Catalog (UHGV)^25^. However, a growing reference database alone does not resolve the fundamental ambiguity of short-read assignment. Many reads classified as viral are not uniquely viral at the sequence level, and sequence homology to other viral genomes or to bacterial and host-derived sequences can increase false positives and compromise abundance estimates^26,27^. At the same time, the large and rapidly growing volume of WMS data has become a major resource for microbiome research^28,29^, yet virome profiling from WMS data remains difficult because of the overwhelming bacterial DNA background. Together, these define both an opportunity and a remaining gap: reference-based virome profiling is now within reach, but it requires methods that can leverage expanding catalogs while maintaining specificity, interpretability, and cross-dataset reproducibility.

Here, we introduce VIP2B (<u>VI</u>rome <u>P</u>rofiler with type <u>IIB</u> restriction sites), a reference-based virome profiling pipeline that performs *in silico* digestion with Type IIB restriction endonucleases to generate short, fixed-length 2B tags for virus detection from WMS data (“DNA-to-2BTag strategy”). Building on our previous work showing that DNA-to-2BTag profiling can mitigate the conventional sensitivity-specificity trade-off in bacteriome profiling^30,31^, VIP2B adapts this paradigm to the virome by generating multi-enzyme 2B tags to improve viral genome coverage and by redesigning the false-positive control model to better accommodate small viral genomes and low viral DNA fractions in metagenomes. Using the latest UHGV as the reference database, VIP2B generates not only species-level taxonomic abundance estimates (i.e., viral operational taxonomic units or vOTUs), but also multiple complementary facets of virome information, including taxonomic coverage, inferred viral gene and gene-cluster profiles, predicted bacterial host profiles, and phenotype-linked signatures. Across diverse taxonomic abundance benchmarking scenarios, VIP2B showed robust performance that was comparable to, and in many cases exceeded, that of state-of-the-art methods. We further show that these multifaceted virome features capture disease-relevant structure beyond abundance alone, improving ordination, differential analysis, and disease classification. Then, by applying VIP2B to 20 public datasets spanning diverse dysbiosis contexts, we demonstrate that bulk metagenomic data harbor substantial recoverable virome information and reveal gut virome patterns associated with health and disease, thereby markedly broadening and enriching the traditionally bacteriome-centered paradigm of microbiome analysis. Together, these results establish VIP2B as a scalable framework for routine, mechanism-oriented virome profiling from metagenomic sequencing data and support the virome as a non-redundant, clinically informative component of the microbiome.

## RESULTS

### Rationale and workflow of VIP2B

Type IIB restriction-modification enzymes cleave DNA on both sides of their recognition site at fixed offsets, generating iso-length fragments (“2B tags”)^32,33^. *In silico* digestion, therefore, provides a compact representation in which each genome is encoded by a characteristic collection of fixed-length 2B tags sampled across many loci, rather than relying on a small set of conserved marker genes or on full-read alignment to reference genomes. We previously demonstrated that the DNA-to-2Btag strategy can simultaneously achieve high sensitivity and high specificity for bacterial, archaeal, and fungal profiling from WMS data^30,31^, substantially mitigating computationally induced false positives that remain a practical limitation for the two conventional strategies (DNA-to-DNA or DND-to-Marker). These results established DNA-to-2Btag as the third profiling paradigm that relaxes the traditional precision-recall trade-off and provides an explicit route to conservative discovery without sacrificing recoverability of true positives. However, extending this concept to the virome is not a simple database swap: viral genomes are short, viral DNA often constitutes a minute fraction of WMS, and viral genomes are highly mosaic, all of which magnify ambiguity and false positives.

To address these challenges, we developed VIP2B, a DNA-to-2BTag workflow for virome profiling (**Fig. 1)**. VIP2B introduces two virome-specific upgrades. First, to increase the yield of informative 2B tags from small viral genomes and to stabilize the coverage of evidence for low-abundance viruses, VIP2B uses a multi-enzyme digestion scheme that pools 2B tags from an eight-enzyme panel (*AlfI*, *BcgI*, *BslFI*, *CjeI*, *CjePI*, *FalI*, *HaeIV*, and *Hin4I*) (**Fig. 1a**). These enzymes were selected because they ranked among the top performers in generating vOTU-specific 2B tags across viral genomes *in silico* digestion and together represented a practical optimum among the 16 Types IIB enzymes evaluated (Supplementary Table 1). Pooling tags from eight enzymes (rather than relying on a single enzyme) expands the vOTU-specific 2B-tag repertoire while preserving a consistent definition of tag-level evidence, thereby increasing the likelihood that true positives contribute multiple independent vOTU-specific tags. Second, VIP2B implements a virome-tailored classifier trained on synthetic virome sequencing data for false-positive recognition. After a first-round alignment against a prebuilt vOTU-specific 2B-tag database derived from UHGV, candidate taxa are evaluated by the classifier, and final abundances are estimated via a second-round alignment to suppress spurious detections while maintaining recall (**Fig. 1b**).

**Figure 1.**
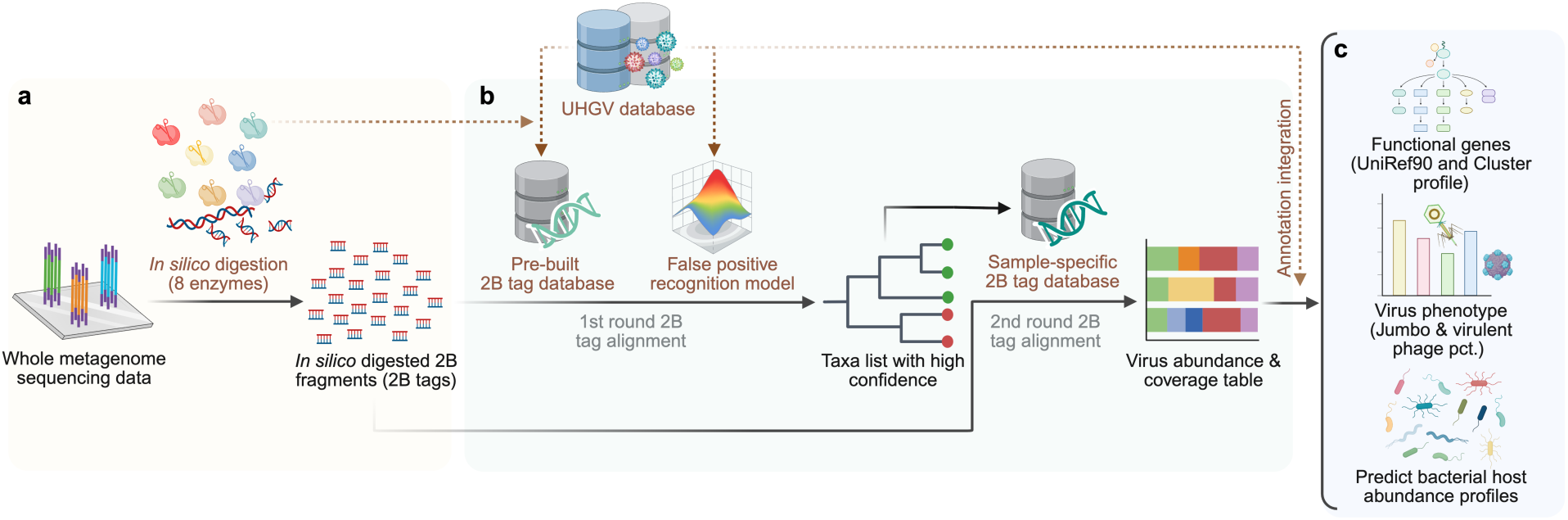
Overview of the VIP2B DNA-to-2Btag workflow for virome profiling from WMS data. (**a**) Conceptual basis of the DNA-to-2Btag strategy: WMS reads, and UHGV viral genomes are projected into a compact 2B tag space by *in silico* 2B enzyme digestion using an eight-enzyme panel (*AlfI*, *BcgI*, *BslFI*, *CjeI*, *CjePI*, *FalI*, *HaeIV,* and *Hin4I*), enabling DNA-to-2Btag profiling as an alternative to conventional DNA-to-marker or DNA-to-DNA approaches. (**b**) A unique two-step reads alignment framework: In the first round of reads alignment, input WMS data are *in silico* digested into 2B-tags. Then, a prebuilt vOTU-specific 2B-tag database is constructed from UHGV and used to identify initial candidates, followed by a false-positive (FP) recognition model that evaluates those candidates from the first-pass screen to infer true viral signal from spurious assignments. In the second round of reads alignment, a sample-specific 2B-tag database is reconstructed from taxa retained after FP recognition to increase the availability of informative vOTU-specific 2B-tags for final quantification. (**c**) Comprehensive downstream readouts are generated by integrating abundance estimates with UHGV metadata, including phenotype-resolved summaries (e.g., jumbo and virulent phage composition), function-resolved profiles (UniRef90 gene family and higher-level functional clusters), predicted bacterial host abundance profiles across taxonomic ranks, and virus abundance summarized across viral taxonomy levels.

In addition, we generate comprehensive downstream readouts by incorporating virus annotations in UHGV, thereby moving beyond the abundance-only outputs that have dominated previous metagenomic profilers, especially existing virome profiling tools. Rather than restricting analysis to taxonomic relative abundance alone, VIP2B produces a multifaceted representation of the virome. These readouts include virus genome coverage (the percentage of the detected vOTU-specific 2b Tags in sequencing data), phenotype-resolved summaries (e.g., jumbo and virulent phage composition), function-resolved profiles (UniRef90 gene families and higher-level functional clusters), predicted bacterial host abundance profiles across taxonomic ranks, and virus abundance summarized at multiple viral taxonomic levels (**Fig. 1c, Methods**).

VIP2B was computationally efficient. Using four CPU threads on a Linux system equipped with an Intel Xeon Platinum 8380 processor, VIP2B profiled a 6.6 GB shotgun metagenomic sample in 6.68 minutes, with a peak memory usage of only 1.09 GB. This short runtime and modest memory requirement support its practical use for large-scale virome profiling of WMS datasets.

### Benchmarking VIP2B for viral taxonomic profiling

To benchmark virome profiling performance in a controlled setting, we generated simulation datasets specifically designed to stress-test common failure modes in virome analysis and used them to compare VIP2B with two representative reference-based profilers, Phanta^24^ and Sylph^34^. To enable a fair head-to-head comparison, we harmonized the reference across profilers by building all three databases from UHGV, thereby standardizing their theoretical detection potential and minimizing confounding from database differences. We implemented five simulation settings (**Fig. 2a**): (1) The standard simulation assessed baseline performance under ideal reference coverage, with both the reference databases and synthetic communities drawn from the full set of available genomes in UHGV, with species richness ranging from 200, 400 to 600 per sample; (2) The dark-matter simulation tested sensitivity to unobserved diversity by building the reference databases from 80% of UHGV genomes while generating synthetic WMS reads from the held-out 20%; (3) The incomplete-genome simulation evaluated robustness to fragmented references by truncating genomes to 70-90% completeness before read simulation; (4) The high-host simulation quantified robustness to overwhelming non-viral background by mixing viral reads with bacterial or (5) human DNA reads at varying host fractions. Each condition included ten replicates. Performance was evaluated in terms of abundance concordance using Bray-Curtis similarity and L2 similarity, and taxon detection using AUPRC, F1 score, precision, and recall.

**Figure 2.**
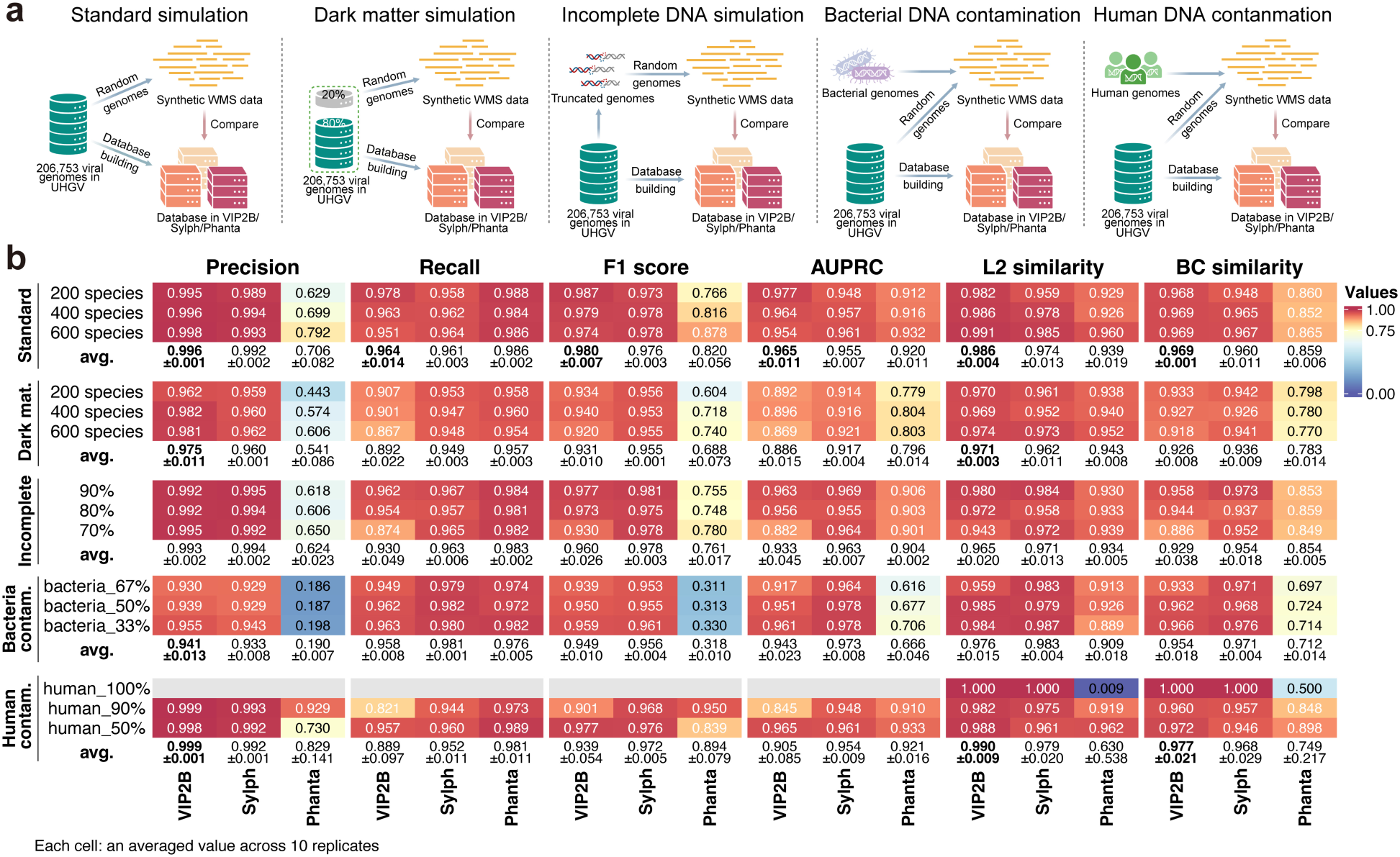
Benchmarking VIP2B across diverse simulation settings. (**a**) Schematic overview of the five simulation settings designed to mimic major challenges in virome profiling from WMS data: standard simulation, dark-matter simulation, incomplete-genome simulation, bacterial DNA contamination, and human DNA contamination. For fair comparison, the reference databases used by VIP2B, Sylph, and Phanta were harmonized by building all three from the same UHGV catalog. (**b**) Heatmap summarizing virome taxonomic profiling performance of VIP2B, Sylph, and Phanta across the five simulation settings. Performance was evaluated using precision, recall, F1 score, and AUPRC for species identification, as well as L2 and BC similarities for abundance estimation. Each cell shows the mean value across 10 replicate simulations for a given condition, and the bottom row within each regime reports the average performance across all tested conditions, together with the corresponding s.d. The averaged metrics where VIP2B achieved the best performance are highlighted in bold.

Across the five benchmarking settings, VIP2B showed consistently high and well-balanced performance, with its clearest advantage in precision and abundance fidelity under challenging backgrounds (**Fig. 2b**). In the standard setting, VIP2B achieved near-perfect precision (average 0.996) while maintaining strong recall (0.964), yielding the highest F1 score (0.980) together with the best concordance to the ground truth by both L2 similarity (0.986) and Bray-Curtis similarity (0.969). This conservative but accurate behavior persisted in the dark-matter and contamination settings, where VIP2B retained markedly higher precision than Phanta and comparable or higher precision than Sylph, while preserving robust recall and abundance agreement. Notably, under bacterial and human DNA contamination, Phanta’s precision deteriorated sharply (average 0.190 and 0.829, respectively), whereas VIP2B remained stable (0.941 and 0.999), indicating substantially greater resistance to spurious viral calls from overwhelming non-viral background. Although Sylph achieved slightly higher recall, and in some settings slightly higher F1 score or AUPRC, this was generally accompanied by reduced precision and L2 similarity relative to VIP2B. Overall, these results indicate that VIP2B provides a favorable operating point for reference-based virome profiling: it preserves strong recoverability of true signals while more effectively suppressing false positives, a property that becomes particularly important for WMS datasets dominated by host or bacterial DNA.

### Taxonomy-guided inference reliably recovers virome functional structure

Beyond taxonomy, the functional repertoire of the virome provides an important layer for interpreting how viruses reshape microbial ecosystems and influence host-associated phenotypes. Yet, direct profiling of viral functional genes from WMS data is intrinsically challenging because many viral genes share sequence homology with bacterial genes, making read-level gene assignment susceptible to ambiguity and false positives. We therefore leveraged the high reliability of VIP2B taxonomic profiling, together with the extensive genome- and gene-level annotations available in UHGV, to infer virome functional composition by projecting vOTU abundance profiles onto curated virus-to-gene relationships encoded in a genome content network (GCN) (**Fig. 3a**, **Methods**).

**Figure 3.**
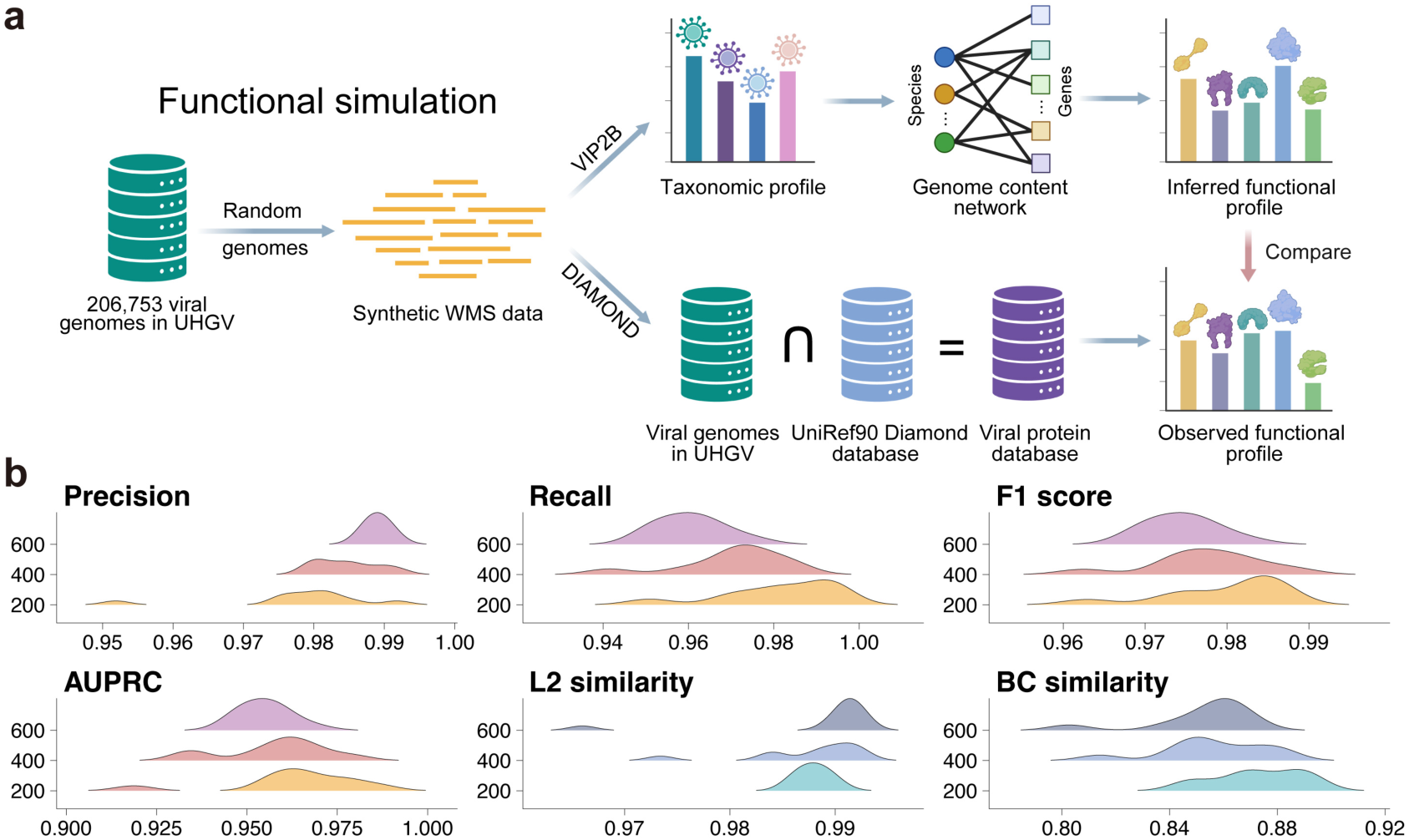
Concordance between VIP2B-predicted and observed functional profiles. (**a**) Schematic overview of the simulation framework used to generate VIP2B-predicted and observed functional profiles. Synthetic whole-metagenome sequencing (WMS) data were generated from randomly selected genomes from UHGV. VIP2B was then used to predict functional profiles by estimating gene abundances from the taxonomic profile through the genome content network (GCN). Observed viral functional profiles were generated by aligning the simulated reads with DIAMOND to a viral protein database built from genes shared between the UHGV viral genomes and the full UniRef90 database. (**b**) Comparison of VIP2B-predicted and observed functional profiles using precision, recall, F1 score, and AUPRC for function detection (red), and L2 and Bray-Curtis similarities for abundance estimation (blue).

We evaluated this strategy in an idealized benchmark containing only viral reads, thereby isolating the fidelity of taxonomy-guided functional inference from bacterial background. VIP2B-predicted gene profiles were compared with observed (direct-mapping-derived) profiles generated from the same simulated datasets (**Methods**). Across species-richness simulations, inferred gene profiles showed very high precision (average 0.984) and strong recall (average 0.971), together with high profile-level concordance by L2 similarity (average 0.988) and Bray-Curtis similarity (average 0.860, **Fig. 3b**). Thus, VIP2B accurately recovered the functional structure of the virome. Additionally, we further summarized gene-level profiles into higher-order gene clusters to reduce sparsity and facilitate downstream interpretation. Together, these results support taxonomy-guided functional inference as a scalable and robust alternative to direct read mapping for virome functional profiling.

### Bulk and VLP metagenomes share more gut viral species than previously recognized

VLP enrichment sequencing and bulk WMS are the two principal strategies for characterizing the human gut virome. Although VLP sequencing is widely used for virome discovery and assembly-based reconstruction, prior studies have suggested that only a limited fraction (10-36%) of viral species detected in VLP data can be recovered from paired bulk WMS data^35–37^. Because these estimates were derived largely from assembly-based workflows, however, they may partly reflect differences in contig recovery and completeness rather than true biological discordance. We therefore re-examined bulk–VLP concordance using an assembly-free, reference-based framework.

We re-analyzed paired bulk and VLP datasets from six human fecal datasets, including two infant datasets and four adult datasets, comprising 2,398 samples from 1,199 bulk-VLP pairs (Supplementary Table 2). Using VIP2B, we directly quantified the extent to which viral species detected in VLP samples were also recovered from the corresponding bulk WMS data. Across datasets, the per-sample overlap was substantially higher than previously reported, with median overlap ranging from 35% to 89% (**Fig. 4a**). When overlap was evaluated at the dataset level by taking the union of species detected across samples within each sequencing strategy, 58-85% of VLP-detected species were also identified in bulk WMS data (**Fig. 4b**). These values markedly exceed earlier assembly-based estimates, indicating that the species-level overlap between bulk and VLP viromes has likely been substantially underestimated.

**Figure 4.**
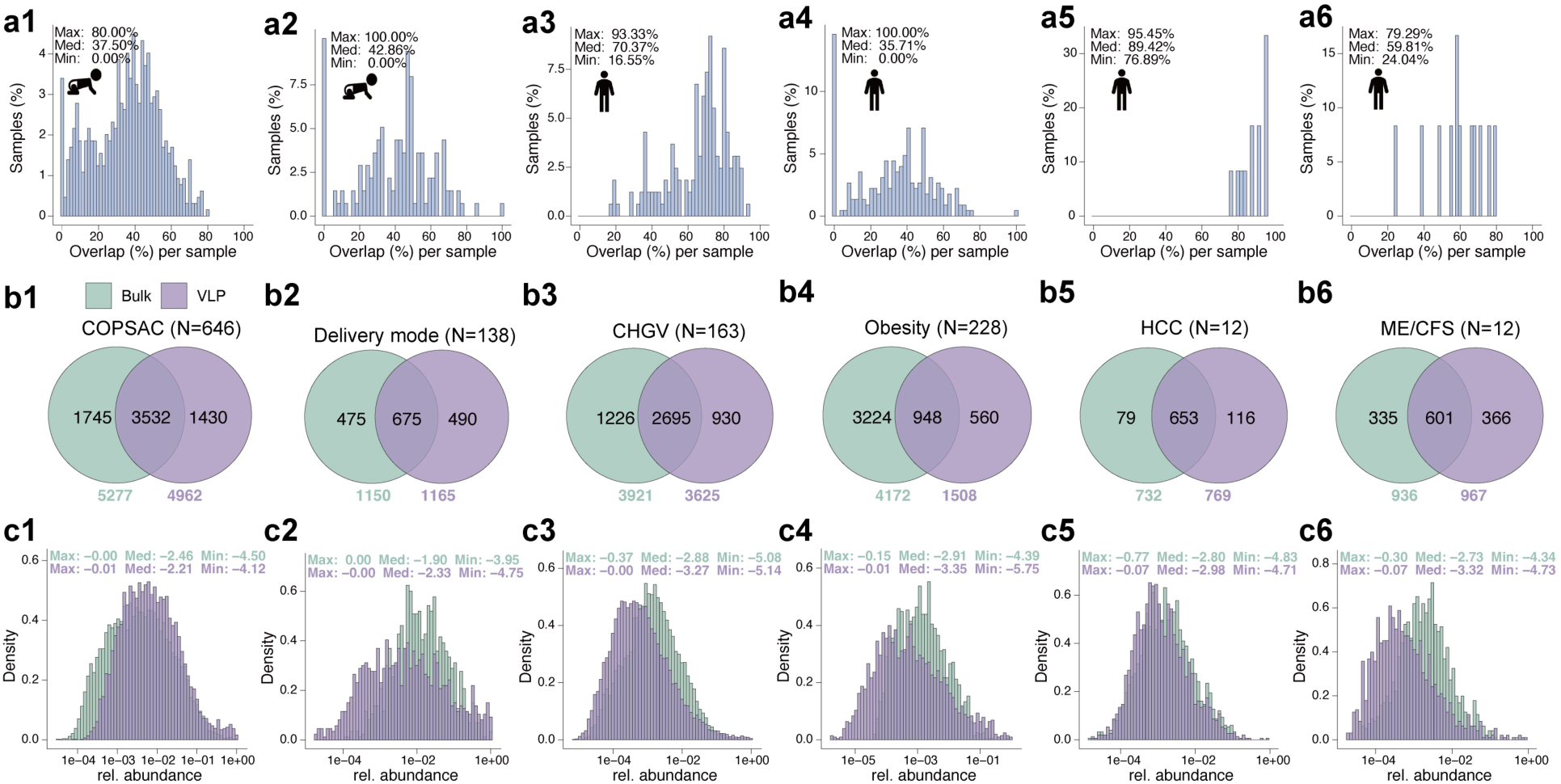
Overlap of viral species between paired bulk shotgun and VLP-enriched metagenomes using VIP2B across six cohorts. Number 1-6 indicate two infant cohorts: COPSAC (Copenhagen Prospective Studies on Asthma in Childhood) and delivery mode, and four adult cohorts: CHGV (Chinese Gut Virome Catalog), obesity, HCC, (hepatocellular carcinoma) and ME/CFS (myalgic encephalomyelitis/chronic fatigue syndrome). (**a1-a6**) Distributions of the percentage of VLP-detected viral species that are also detected in the paired bulk metagenome for each sample. Minimum (Min), median (Med), and maximum (Max) of the percentages are annotated. (**b1-b6**) Venn diagrams showing the numbers of shared and method-specific viral species recovered from bulk and VLP data within each cohort. (**c1-c6**) Density distributions of cumulative abundances of overlapping viral species in bulk and VLP data. Minimum (Min), median (Med), and maximum (Max) of the log10-scaled abundances are annotated.

Despite this higher-than-expected overlap, the shared fraction was not uniform across populations. Infant datasets showed consistently lower overlap than adult datasets, with a median per-sample overlap of 37.5% in the Copenhagen Prospective Studies on Asthma in Childhood (COPSAC) dataset and 42.9% in the delivery-mode dataset, whereas three adult datasets showed median overlap values near or above 60%. This pattern is consistent with the greater interpersonal variability and developmental instability of the infant gut virome reported previously, which would be expected to reduce cross-method recoverability at the species level^38,39^. At the same time, a substantial fraction of the detected species remained sequencing-method-specific across all datasets. Notably, in the largest datasets, bulk WMS tended to recover more phage species overall than VLP sequencing, suggesting that at sufficient scale, bulk WMS data can capture not only free viral particles but also a broader spectrum of host-associated viral signals.

We next asked whether the species shared between bulk and VLP data were represented similarly in abundance. Despite the substantially larger species-level overlap, abundance distributions for the shared species differed markedly between bulk and VLP data in every dataset (two-sample Kolmogorov-Smirnov test, P < 4.16×10^−9^; **Fig. 4c** and Supplementary Table 3). Similar results were obtained with Sylph-derived profiles (Supplementary Fig. 1). Together, these findings indicate that bulk WMS contains substantially more recoverable virome information than is commonly assumed and that VLP enrichment systematically reshapes the quantitative structure of the observed virome.

### Multifaceted virome profiling increases sensitivity to gut dysbiosis

To assess the extent to which bulk WMS can support biologically informative virome analysis at scale, and to test whether VIP2B’s expanded feature space improves sensitivity for detecting disease-associated virome alterations, we analyzed 3,685 samples from 20 publicly available gut WMS datasets covering 16 homeostatic and dysbiosis states (Supplementary Table 2). We first asked whether case-control differences could be detected at the community level using distinct classes of VIP2B-derived virome features. Across the 20 datasets, at least one VIP2B-derived profile showed significant case-control separation by PERMANOVA in 17 datasets (P < 0.05; **Fig. 5a**). Two hepatocellular carcinoma (HCC)-related datasets showed the strongest effects, with disease status explaining more than 20% of the variance in virome composition (R^2^ > 0.2, Supplementary Fig. 2). Across datasets, VIP2B’s multifaceted profiles provided more sensitive detection of dysbiosis-associated virome structure than abundance-only profiling. Specifically, in 19 of 20 datasets, at least one VIP2B-derived feature type yielded a smaller or equal PERMANOVA P value together with a larger F statistic than Sylph-derived taxonomic abundance profiles (**Fig. 5a, 5b**). This advantage was particularly evident in datasets such as multiple sclerosis (MS) and colorectal cancer (CRC), in which Sylph-derived abundance profiles did not reach significance, whereas several VIP2B-derived profiles (including taxonomic abundance, taxonomic coverage, gene profiles, and gene-cluster profiles) did achieve statistical significance. Among all VIP2B features, taxonomic coverage was the single best-performing profile in 13 of 20 datasets, indicating that dysbiosis-associated virome differences are often more sensitively captured by the breadth of viral genome detection than by relative abundance alone.

**Figure 5.**
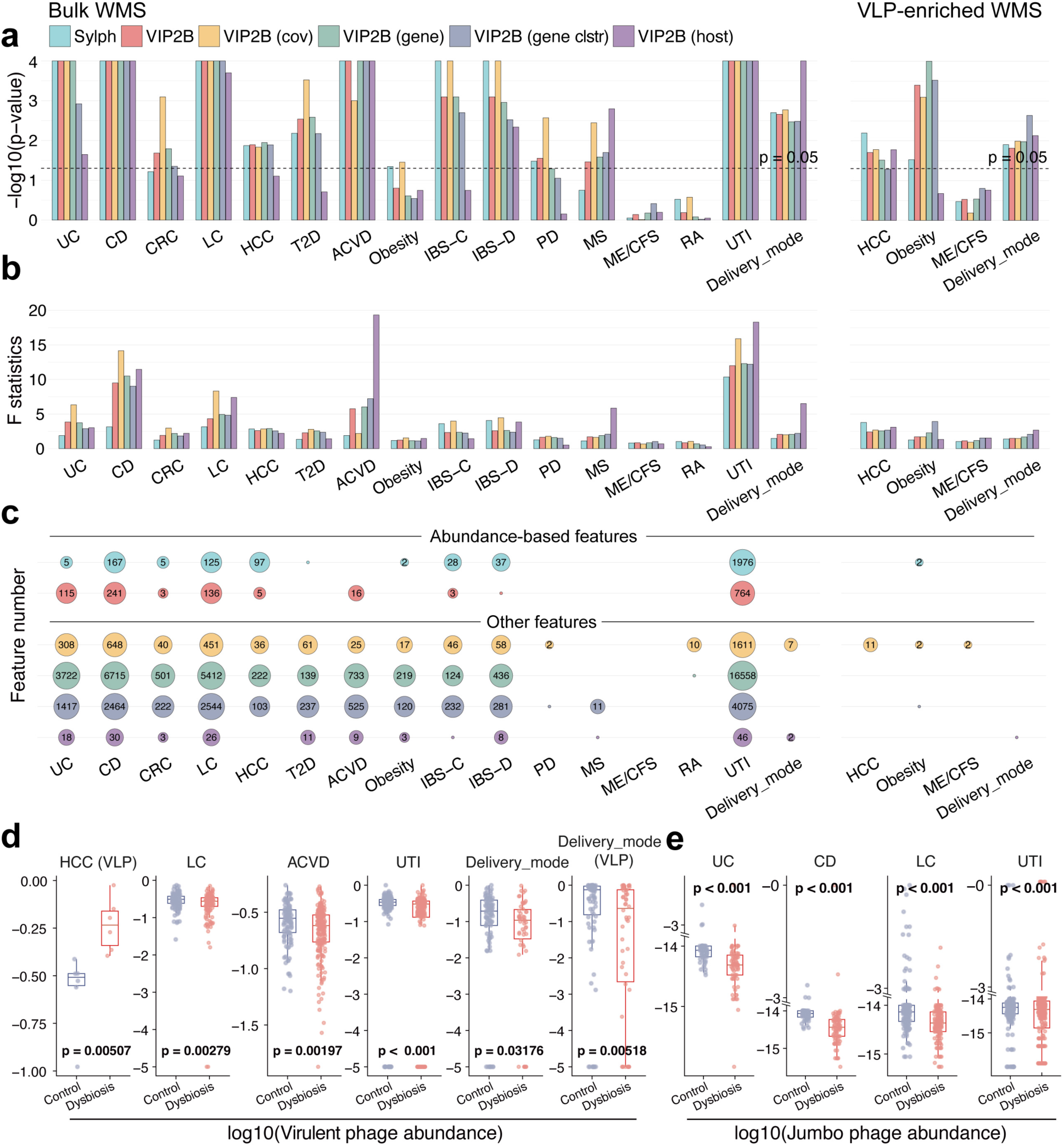
Multi-faceted virome profiling reveals dysbiosis-associated structure beyond taxonomic abundance across 20 public gut metagenomic datasets. (**a-b**) PERMANOVA (9,999 permutations) testing case-control differences in virome beta diversity across 20 datasets using Sylph-derived taxonomic abundance profiles and multiple VIP2B-derived feature classes. Results are shown as (**a**) -log10(P) values and (**b**) F statistics. VIP2B-derived profiles include viral taxonomic abundance (vOTU level), taxonomic coverage, functional profiles (gene and gene-cluster levels), and predicted host taxonomic profiles. Results for additional taxonomic levels are shown in Supplementary Fig. 8. (**c**) Numbers of dysbiosis-associated viral features identified by MaAsLin3 (FDR-adjusted P < 0.25) across feature classes and datasets. (**d-e**) Boxplots showing disease datasets in which (**d**) virulent-phage abundance and (**e**) jumbo-phage abundance differed significantly between controls and dysbiosis. P values were calculated using two-sided Wilcoxon rank-sum tests. Clstr: Cluster; LC, liver cirrhosis; UTI, urinary tract infection; ACVD, atherosclerotic cardiovascular disease; HCC, hepatocellular carcinoma; VLP, virus-like particle-enriched metagenomics. The delivery-mode dataset comprises infant stool samples stratified by vaginal versus Cesarean delivery.

We next performed differential feature analysis to identify disease-associated viral signals at finer resolution (**Methods**). At least one VIP2B-derived non-abundance feature type identified more differential signals than taxonomic abundance profiles from either Sylph or VIP2B in 18 of 20 datasets (**Fig. 5c**). Notably, taxonomic coverage identified dysbiosis-associated viral species in five additional datasets in which abundance-based features were less sensitive, further supporting the value of coverage as a complementary descriptor of virome perturbation. Functional profiles added another layer of resolution: gene- and gene-cluster-level features uncovered substantially more differential signals in five datasets, including CRC, Atherosclerotic Cardiovascular Disease (ACVD), obesity, and MS, than either abundance- or coverage-based features. To assess whether these additional functional signals were biologically informative, we examined disease-associated phage gene clusters in an inflammatory bowel disease (IBD) dataset^40^. Across both Crohn’s disease (CD) and ulcerative colitis (UC), VIP2B consistently detected depletion of phage gene clusters linked to host adaptation, including diversity-generating retroelements (DGRs), as well as features related to stable prophage maintenance, including ParB-associated partition systems (Supplementary Fig. 3). These functional changes are concordant with prior evidence for marked virome disruption in IBD and suggest that phage-associated adaptive capacity and maintenance programs are perturbed in inflammatory disease states^9,10,41–43^.

We further incorporated phenotype-linked virome summaries into VIP2B by quantifying virulent- and lysogenic-associated phage fractions and jumbo phage abundance. These features were motivated by growing evidence that phage lifestyle can influence microbiome instability^4,44^, and that jumbo phages, although prevalent in the gut, remain underexplored in the context of human disease^45^. Across multiple datasets, these higher-order virome summaries revealed significant case-control differences in several disease settings (**Fig. 5d, 5e**). Virulent-phage abundance differed significantly in the VLP-based HCC dataset (P = 0.005), liver cirrhosis (LC; P = 0.003), ACVD (P = 0.002), urinary tract infection (UTI, P < 0.001), and the delivery-mode dataset in both bulk WMS (P = 0.032) and VLP data (P = 0.005). Jumbo-phage abundance was also significantly altered in UC (P < 0.001), CD (P < 0.001), LC (P < 0.001), and UTI (P < 0.001). Collectively, these analyses show that extending virome profiling beyond taxonomic abundance to include coverage, functional, host-linked, and phenotype-resolved summaries increases sensitivity for detecting gut dysbiosis across heterogeneous datasets and provides a richer framework for linking virome variation recovered from bulk WMS data to disease biology.

### Bacteriome-virome alpha-diversity coupling does not generalize as a marker of dysbiosis

Despite the long-standing dominance of bacteriome-centered microbiome analyses, we hypothesize that the virome provides non-redundant ecological and disease-relevant information that is not fully captured by bacterial composition alone. Rather than serving merely as a bystander or passive reflection of bacteriome shifts, the virome represents a complementary dimension of microbiome organization in its own right. Against this backdrop, a recent meta-analysis proposed that, relative to healthy controls, the correlation between bacteriome and virome alpha diversity is weakened in dysbiosis^46^. We therefore sought to test this hypothesis systematically across the same 20 datasets analyzed above. Within each dataset, we quantified the association between bacterial and viral Shannon diversity and compared the strength of these correlations between controls and dysbiosis (**Methods**).

Contrary to the proposed general trend, we did not observe a consistent reduction in bacteriome-virome alpha-diversity coupling in dysbiosis when virome profiles were derived from bulk WMS data. Across the 16 disease or dysbiosis contexts examined, the direction and magnitude of change varied substantially across datasets, and only seven showed higher bacteriome-virome correlations in controls than in dysbiosis. At the aggregate level, the difference in correlation strength between healthy and diseased states was not statistically significant across the 16 conditions (**Fig. 6a**). We next repeated the analysis in the four datasets with paired VLP data and again found no significant decrease in Shannon-index correlations in dysbiosis (**Fig. 6b**). Because VLP enrichment preferentially captures free viral particles and is often enriched for virulent phages, we further asked whether a more specific ecology-informed coupling metric might be more informative. We therefore examined the correlation between bacterial alpha diversity and virulent-phage alpha diversity inferred from bulk WMS data but again found no consistent case-control difference across datasets (**Fig. 6c**). These conclusions were unchanged when Simpson diversity was used in place of Shannon diversity (Supplementary Fig. 4a) or when the analysis was repeated with Sylph-derived virome profiles (Supplementary Fig. 4b, 4c).

**Figure 6.**
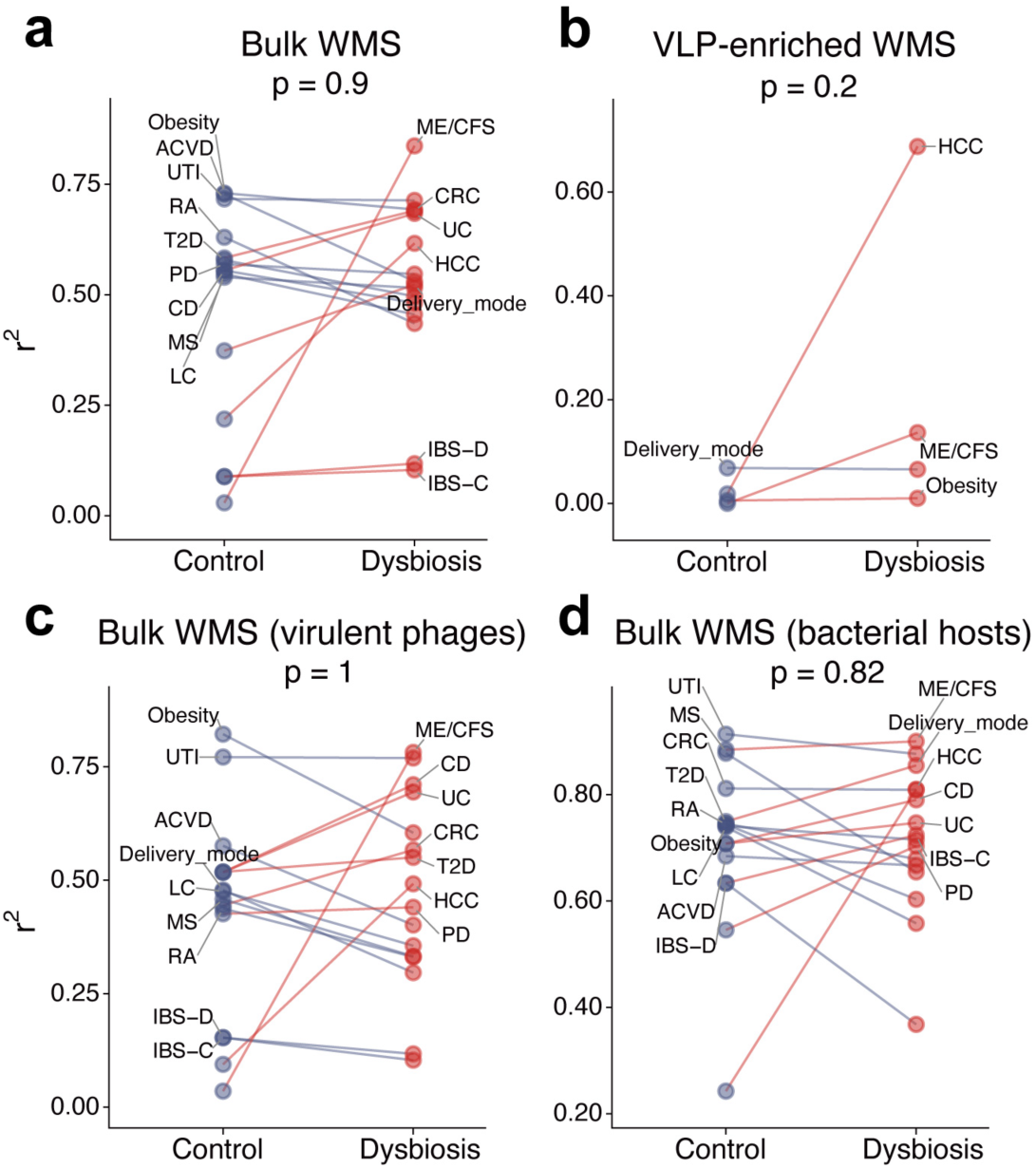
Cross-kingdom alpha-diversity coupling is not consistently weakened in dysbiosis. Pearson correlation coefficients of determination (r^2^) comparing bacteriome–virome Shannon-diversity coupling between controls and dysbiosis across datasets. Viral profiles were generated by VIP2B at the species level. Analyses are shown for (**a**) 16 bulk WMS datasets using the full virome, (**b**) 4 VLP enrichment metagenomic datasets, (**c**) the same 16 bulk WMS datasets with viral diversity restricted to virulent phages, and (**d**) the same 16 bulk WMS datasets with bacterial diversity restricted to the predicted hosts of the detected viruses. Each connected pair denotes one dataset. Blue lines indicate stronger coupling in controls, whereas red lines indicate stronger coupling in dysbiosis; labels are positioned on the side with the higher r^2^ value. P values shown in the above panels were obtained from paired comparisons across datasets.

We additionally considered a more targeted hypothesis: if cross-kingdom coupling were primarily driven by viruses that infect bacteria, then the relationship might become more apparent when analysis was restricted to bacterium-associated viral diversity rather than the virome as a whole. Leveraging VIP2B-predicted bacterial host profiles, we therefore tested the association between viral alpha diversity and the alpha diversity of their predicted bacterial hosts. This analysis likewise showed no systematic difference between controls and dysbiosis (P = 0.2; **Fig. 6d**), indicating that even when restricted to host-linked viral diversity, alpha-diversity coupling does not emerge as a robust marker of dysbiosis. These findings support the view that the virome contributes a partially independent layer of information and that virus–bacterium relationships in dysbiosis are more complex than a single diversity-coupling statistic can capture.

### Viral features contribute non-redundant predictive information beyond bacteriome-centered models

Because cross-kingdom alpha-diversity coupling did not generalize as a robust marker of dysbiosis, we next asked whether the virome contributes more directly useful predictive information for disease classification. Given that VIP2B-derived virome profiles showed strong and often superior case-control separation across multiple feature layers (**Fig. 3**), we tested whether viral features, either alone or in combination with bacterial profiles, could improve classification performance beyond bacteriome-centered models. To address this, we selected 15 from the 20 datasets analyzed above that contained at least 50 samples each and benchmarked two representative classifiers: Elastic Net (EN) and Random Forest (RF). Model performance was evaluated using five repeats of 10-fold cross-validation, and predictive accuracy was summarized by the mean area under the receiver operating characteristic curve (AUROC) across folds and repeats (**Methods**). We compared six feature sets: bacterial taxonomic profiles generated by MAP2B across all taxonomic ranks (MAP2B taxa), viral taxonomic profiles generated by VIP2B across all taxonomic ranks (VIP2B taxa), viral genome coverage profiles (VIP2B cov), viral functional features including gene and gene-cluster profiles (VIP2B func), the full VIP2B feature panel integrating taxonomic, coverage, functional, predicted host, virulent phage and jumbo phage features (VIP2B all), and a combined feature space containing all bacterial and viral features (All combined).

Across datasets, viral features frequently matched or exceeded the predictive performance of bacteriome-centered models. Under EN modeling, at least one VIP2B-derived feature set, either alone or in combination with bacterial profiles, outperformed the bacterial baseline in 11 of 15 datasets (73%; **Fig. 7a**). Under RF modeling, the corresponding proportion was 9 of 15 datasets (60%; **Fig. 7b**). Overall, across the 30 dataset-classifier evaluations, incorporating VIP2B-derived virome features improved AUROC relative to bacteria-only models in 20 cases (67%). This added value was most apparent in settings where virome disruption was likely to be biologically informative. In CRC, for example, viral features consistently outperformed the bacterial features under both classifiers, increasing AUROC from 0.610 to 0.752 with EN and from 0.666 to 0.758 with RF. Likewise, in the paired VLP delivery-mode dataset, integrating virome information markedly improved prediction, with AUROC increasing from 0.509 to 0.684 under EN and from 0.492 to 0.673 under RF. Viral features also substantially improved prediction in the obesity dataset, where AUROC rose from 0.670 to 0.720 under RF. Together, these results indicate that the virome can provide practically meaningful predictive signals in a substantial subset of dysbiosis contexts and support moving microbiome-based classification frameworks beyond bacteriome-centered designs toward models that explicitly incorporate the virome as an independent axis of disease-relevant variation.

**Figure 7.**
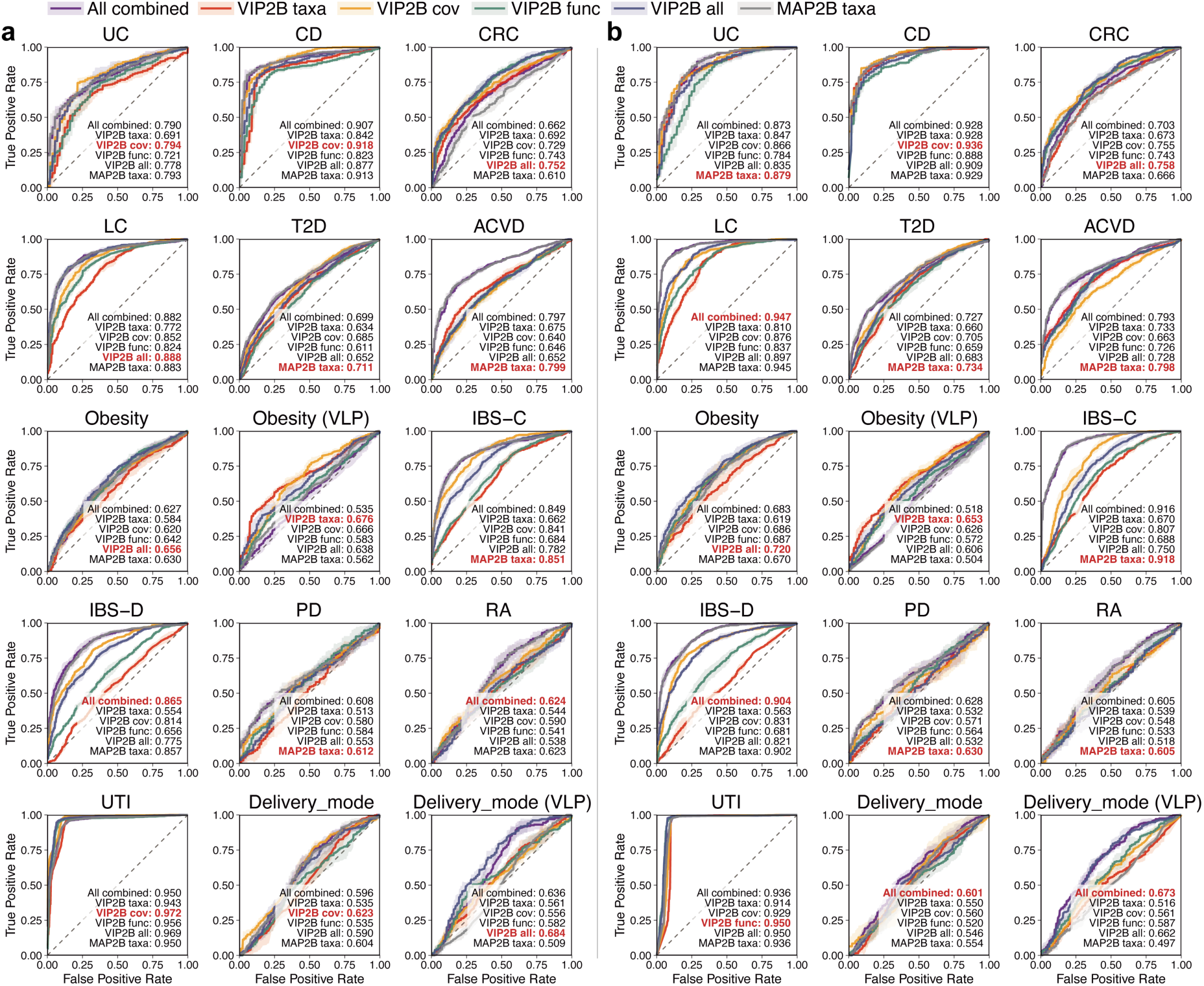
Disease-classification performance of bacterial, viral, and integrated microbiome feature sets across 15 datasets. (**a-b**) Receiver operating characteristic (ROC) curves showing disease-classification performance across 15 datasets using (**a**) Elastic Net and (**b**) Random Forest models. Curves represent the mean performance across five repeats of 10-fold cross-validation, and shaded regions denote 95% confidence intervals. Mean area under the ROC curve (AUROC) values are shown for each feature set, with the best-performing model in each dataset highlighted in red. Feature sets include MAP2B taxa (bacterial taxonomic abundances across all taxonomic ranks), VIP2B taxa (viral taxonomic abundances across taxonomic ranks), VIP2B cov (viral genome coverage/completeness), VIP2B func (viral functional features, combining gene- and gene-cluster-level profiles), VIP2B all (the full VIP2B feature panel, including taxonomic, coverage, functional, predicted host, virulent-phage and jumbo-phage accumulated abundances), and All combined (the union of all MAP2B and VIP2B features). TPR, true positive rate; FPR, false positive rate.

### Large-scale gut virome profiling reveals two reproducible community types associated with health

Having shown that the virome contributes non-redundant ecological and predictive information beyond bacteriome-centered analyses, we finally asked whether large-scale standardized virome profiling could reveal organizational patterns of the human gut virome itself. In particular, we sought to determine whether reproducible gut virome community types exist across studies and sequencing strategies, and whether such community states are linked to host health. To address this, we aggregated 6,090 adult stool metagenomes from 18 studies, including both bulk and VLP datasets (Supplementary Table 2). We focused on taxonomic coverage profiles, which showed strong and consistent performance in both compositional analyses and disease classification across the preceding sections (**Fig. 3** and **Fig. 5**).

We first assessed whether the data supported discrete virome community structure. Samples were clustered using partitioning around medoids (PAM) across three distance measures, while varying the number of clusters from K = 2 to K = 10. Clustering quality was evaluated using the Calinski-Harabasz (CH) index^47^, prediction strength^48^, and Silhouette score^49^, which collectively assess cluster compactness and separation. Across distance metrics, all three criteria consistently favored K = 2, with maximal scores at K = 2 (**Fig. 8a-c**). However, prediction strength values below 0.8 and silhouette scores below 0.2 indicated weak overall support for well-separated clusters (**Fig. 8b-c**)^50^. We further tested whether the data support no clustering (K = 1) by fitting Dirichlet multinomial mixture (DMM) models and comparing Bayesian information criterion (BIC) values^51^. BIC decreased sharply from K = 1 to K = 2, arguing against a single cluster model, but continued to slowly decrease up to K = 10 without a clear minimum (Supplementary Fig. 5a), consistent with diminishing returns as additional clusters are added. These analyses support the presence of separated community structure, with K = 2 representing the most parsimonious summary of the data.

**Figure 8.**
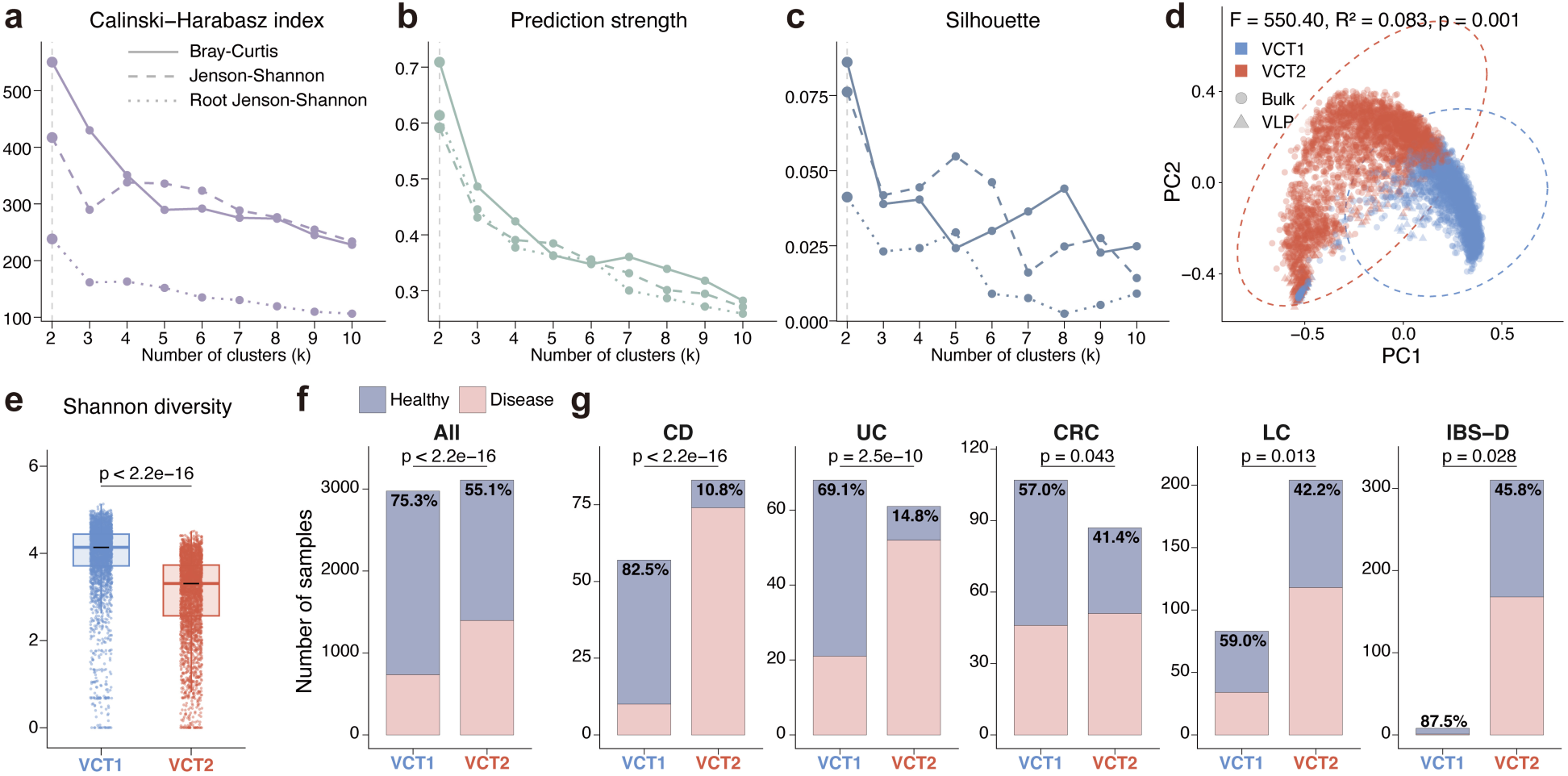
VIP2B coverage profiles from 6,090 stool metagenomes suggest two putative virome community types. (**a-c**) Cluster validity metrics for partitioning around medoids (PAM) clustering of species-level VIP2B taxonomic coverage profiles from 6,090 stool metagenomes across K = 2-10 clusters, evaluated using (**a**) the Calinski-Harabasz index, (**b**) prediction strength, and (**c**) silhouette score. Clustering was performed using Bray-Curtis, Jensen-Shannon, and root Jensen-Shannon distances. (**d**) Principal coordinates analysis (PCoA) of coverage profiles, colored by the two putative virome community types (VCT1, blue; VCT2, red). Sample shape indicates sequencing strategy (circles, bulk WMS data; triangles, VLP sequencing data). Group separation was assessed by PERMANOVA (999 permutations). (**e**) Viral alpha diversity, measured by the Shannon index, in VCT1 and VCT2. P value was calculated using a two-sided Wilcoxon rank-sum test. (**f-g**) Distribution of healthy and disease samples across (**f**) the full aggregated dataset and (**g**) individual disease datasets. Numbers within bars indicate the percentage of healthy samples in each virome community type. P values were calculated using two-sided Fisher’s exact tests. Disease datasets include Crohn’s disease (CD), ulcerative colitis (UC), colorectal cancer (CRC), liver cirrhosis (LC), and irritable bowel syndrome with diarrhea (IBS-D).

Consistent with this analysis, principal coordinates analysis (PCoA) also showed separation (PERMANOVA test, p < 0.05), revealing two partially overlapping groups that we refer to as the putative virome community types, VCT1 and VCT2 (**Fig. 8d**). Despite the limited separation between VCT1 and VCT2, we observed reproducible biological differences between the groups. VCT1 exhibited significantly higher viral alpha diversity than VCT2, as measured by the Shannon index (**Fig. 8e**). Moreover, healthy samples were significantly enriched in VCT1 across the full collection (**Fig. 8f**). This enrichment was also observed within multiple disease datasets, including CD, UC, CRC, LC, and IBS-D (**Fig. 8g**), consistent with prior reports linking virome enterotypes to disease contexts^13,52–54^.

We next asked whether this two-group structure was specific to coverage-based virome representation or whether it recurred across other VIP2B-derived feature layers. Using species-level taxonomic abundance and gene-level functional profiles, we again observed evidence for a two-group organization. In both cases, the CH index and prediction strength supported K = 2, although silhouette scores remained low and DMM-based BIC values again failed to identify a sharp optimum (Supplementary Figs. 5b, 5c, 6a-d and 7a-d). Thus, the same overall picture emerged across feature spaces: a reproducible but weakly separated two-group structure rather than a set of strongly discrete community types. Consistent with the coverage-based analysis, both abundance- and function-derived groupings identified one group with higher viral alpha diversity and a greater proportion of healthy samples (Supplementary Figs. 6e and 7e). When stratified by disease, enrichment of healthy samples based on abundance and functional profiles was detected in three and two disease datasets, respectively, and these always included CD (Supplementary Figs. 6f-g and 7f-g). Nevertheless, the two-cluster assignments derived from coverage, abundance, and functional profiles showed moderate-to-strong concordance (odds ratios ≥ 4.8, p ≤ 5.21 × 10^−188^, Supplementary Fig. 6h and 7h-i). These results suggest that rather than resolving into strongly separated enterotype-like classes, the human gut virome appears to organize into two broad and weakly separated but reproducible community states. One of these states, VCT1, is consistently associated with higher viral alpha diversity and enrichment of healthy samples, suggesting a potentially more favorable ecological configuration of the gut virome.

## DISCUSSION

The gut virome has remained less tractable than the bacteriome, not because it is biologically less important, but because it has been substantially harder to profile reproducibly and at scale across studies. This technical asymmetry has reinforced a bacteriome-centered view of the microbiome, despite the central ecological role of phages in shaping bacterial abundance, evolution, and host-associated community function. Here, we developed VIP2B to help close this gap. By extending the DNA-to-2BTag framework to the virome, VIP2B provides an assembly-free, reference-based approach for conservative and scalable virome profiling directly from whole-metagenome sequencing data.

A key advance of VIP2B is its ability to maintain specificity in settings where reference-based virome profiling is especially vulnerable to false positives, including small viral genomes, incomplete references, and overwhelming bacterial or host DNA background. Across simulation settings designed to mimic these challenges, VIP2B showed strong concordance in abundance estimates and high detection accuracy while remaining robust to bacterial and human DNA contamination. These properties are particularly important for bulk WMS, where even modest false-positive rates can distort ecological and clinical interpretation. Our analyses also revise the prevailing view of how much virome information can be recovered from bulk metagenomic data. Previous assembly-based comparisons have generally implied that bulk WMS captures only a limited subset of the gut virome observed by VLP-enriched sequencing. In contrast, using a standardized, assembly-free framework, we found substantially greater overlap in the viral species detected between paired bulk and VLP datasets. At the same time, the two sequencing strategies remained systematically different in their abundance structures and overall community compositions. Thus, bulk WMS is not merely a poor proxy for the virome; rather, it contains a substantial recoverable viral signal that has likely been underestimated by assembly-centric workflows, whereas VLP enrichment remains valuable for reshaping quantitative representation and enriching free-virion signal.

Beyond taxonomy, VIP2B was designed to recover additional layers of virome structure, including taxonomic coverage, inferred functional profiles, predicted host associations, and phenotype-linked summaries. These features repeatedly improved sensitivity for detecting dysbiosis-associated structure and, in many settings, provided stronger discriminatory power than abundance alone. This suggests that disease-associated virome remodeling is intrinsically multi-dimensional. Relative abundance captures only one aspect of viral community perturbation, whereas coverage-, function-, and lifestyle-linked summaries reveal additional axes of ecological change. In this sense, VIP2B shifts virome analysis from a largely descriptive cataloging exercise toward a richer representation of viral ecological and functional organization. This broader feature space also changes how the virome should be positioned within microbiome research. Viral features repeatedly provided non-redundant information beyond bacterial taxonomic profiles and frequently improved disease classification, either alone or in combination with bacterial features. The implication is not simply that viruses mirror bacterial changes, but that they capture additional aspects of ecosystem disruption that cannot be recovered from bacteriome-only models. More broadly, these findings argue against a strictly bacteria-centered view of the microbiome and instead support an integrated framework in which viruses and bacteria are treated as complementary layers of host-associated ecosystem biology.

Our results also caution against overly simple ecological summaries of virome dysbiosis. A recently proposed weakening of bacteriome-virome alpha-diversity coupling in disease did not generalize across the heterogeneous disease contexts analyzed here. Instead, virome-associated signals were feature-specific and context-dependent, indicating that clinically relevant virome perturbation is unlikely to be captured by a single cross-kingdom diversity metric. At a larger scale, standardized profiling of 6,090 adult stool metagenomes further suggested that the gut virome may organize into two broad but weakly separated community states, one of which was reproducibly associated with higher viral alpha diversity and with enrichment in healthy samples. Although these groups do not support sharply discrete enterotype-like classes, they suggest that the gut virome may nonetheless adopt higher-order ecological configurations relevant to host health.

Several limitations should be acknowledged. VIP2B remains reference-informed and therefore depends on the completeness and curation of available viral genome catalogs. Assembly-free profiling complements rather than replaces *de novo* assembly, which remains indispensable for novel virus discovery and for improving reference resources. In addition, our function-resolved analyses are intentionally conservative and prioritize reliable sample-level structure over exhaustive gene recovery. Future work should focus on improving reference breadth, optimizing 2B tags selection and enzyme design, and more deeply integrating viral and bacterial layers from the same bulk WMS data to strengthen inference of multi-kingdom ecology. In summary, VIP2B provides a scalable and conservative approach to virome profiling from metagenomic data, extending analysis beyond taxonomy to include coverage, functional, host-linked, and lifestyle-related signals. By enabling standardized virome profiling across datasets and sequencing strategies, VIP2B shows that the virome is not merely an accessory readout of the microbiome, but a non-redundant and clinically informative layer of host-associated biology.

## METHODS

### Overview of VIP2B

VIP2B is a DNA-to-2Btag virome profiling workflow for whole-metagenome sequencing (WMS) data. The pipeline operates in two stages. In the first stage, WMS reads are converted into Type IIB restriction 2B tags via *in silico* digestion and mapped to a precomputed virus-specific unique-tag index to generate candidate detections together with diagnostic summary statistics. Candidate taxa are then passed through a conservative false-positive recognition model to predict a set of true-positive taxa. In the second stage, VIP2B reconstructs a sample-specific unique-tag database restricted to the predicted true-positive taxa, thereby increasing the number of informative tags per taxon and enabling robust relative abundance estimation from a second round of tag mapping.

### Construction of the virus 2Btag reference database

We constructed a virus 2B tag reference database from the UHGV genome collection. For each reference unit 𝑖 (e.g., a species-level viral cluster/vOTU; hereafter “taxon”), we performed *in silico* digestion of all associated genomes using a specified set of Type IIB restriction enzymes. Digestion was implemented by scanning both forward and reverse-complement strands using enzyme-specific recognition patterns (regular-expression-based matching) and enumerating all resulting fixed-length tags. 2B tags were stored in a compact key–value index using marisa-trie, where each entry maps a tag sequence to its originating genome identifier and observed within-genome copy number (encoded as genomeID + copy number). To scale to large reference collections, the index was partitioned into multiple sub-databases, each containing a fixed number of genomes. To enable high-specificity detection, VIP2B uses a pre-constructed unique 2B tag database, comprising 2B tags that are specific to a taxon relative to all other taxa in the UHGV database reference. Unique 2B tags were derived by comparing the set of theoretically digestible tags from each taxon to all other taxa at a given taxonomy resolution. The database construction module outputs (*i*) a Marisa index for tag lookup and (*ii*) a companion statistics table recording theoretical tag counts per genome and per taxon.

### *In silico* digestion of WMS reads into 2B tags

Given raw WMS reads, VIP2B extracts *in silico* 2B tags by applying the same enzyme recognition patterns used in database construction. Digested tags are written as a compressed FASTA file (*.fa.gz), together with sample-level digestion summary statistics (e.g., number of tags recovered and tag-to-read ratio). Unless otherwise stated, digestion scans both strands and retains exact tag sequences as the matching units for downstream mapping. To increase the yield of informative tags for short viral genomes and to improve robustness under low virome fractions in WMS, VIP2B uses a multi-enzyme Type IIB digestion scheme. By default, VIP2B employs an 8-enzyme panel (*AlfI*, *BcgI*, *BslFI*, *CjeI*, *CjePI*, *FalI*, *HaeIV*, and *Hin4I*). For each read, VIP2B scans both strands against the selected enzyme recognition patterns and aggregates all resulting fixed length 2B tags into a single pooled tag set (union pooling across enzymes). In addition, an embedded human genome (GRCh38.p14 and CHM13v2.0) has been used to remove any possible host DNA contamination, so no additional pre-host removal steps are required. For workflows requiring explicit removal of other host-derived tags, VIP2B can generate a host tag index from one or more host genomes and exclude tags matching the host index prior to microbial mapping.

### First-round mapping and computation of diagnostic features

Let 𝑖 index a taxon in the virus reference database. Denote by 𝐻_i_the total number of 2B tags theoretically generated by *in silico* digestion of taxon 𝑖’s genome(s), and by 𝐸_i_the number of single-copy, taxon-specific unique tags for taxon 𝑖 with respect to all other taxa in the database. For a given WMS sample, VIP2B maps digested tags to the preconstructed unique 2B tag database and collects 𝑄_i_ (the total count of mapped unique tags assigned to taxon 𝑖) and 𝑈_i_(the number of distinct unique tags observed for taxon 𝑖, namely nonredundant unique tags). We define the genome coverage of taxon 𝑖 as 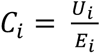. Following the coverage correction used in MAP2B^31^, we infer the number of sequenced unique tags as 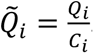. We further compute: (*i*) the taxonomic count: 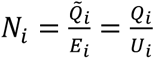, (*ii*) the sequence count: 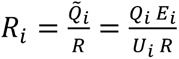, where 𝑅 is the total number of reads in the sample, and (*iii*) a G-score summarizing joint support from breadth and depth: 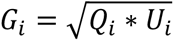. These features are log-transformed prior to false-positive recognition.

### False-positive recognition

VIP2B supports two complementary false-positive (FP) control modes: (1) Threshold-based filtering. As a conservative default, VIP2B can label taxa as present/absent by applying a minimum G-score cutoff 𝐺_i_ > 𝑇, which 𝑇 is user-configurable. (2) Machine-learning FP recognition. VIP2B also implements a supervised FP recognition model (XGBoost), which takes as input the log-transformed feature vector: log (𝐶_i_), log (𝑅_i_), log (𝑁_i_), and log (𝐺_i_). The trained XGBoost model outputs a probability 𝑝_i_that taxon 𝑖 represents a true-positive detection. VIP2B reports taxa as present when 𝑝_i_ > γ, where γ is user-configurable (default γ = 0.5). This FP recognition step is designed to be conservative under low-signal virome conditions and is applied before sample-specific database reconstruction and final quantification. A total of 50 simulated samples were used to train the XGBoost model. These samples covered communities containing 100, 200, …, 1,000 species, with five replicates generated for each species richness level, following the same simulation strategy used for systematic benchmarking. Different simulation seeds were used to ensure that the training samples were distinct from all benchmarking samples.

### Sample-specific database reconstruction

To improve quantification accuracy after FP removal, VIP2B reconstructs a sample-specific unique-tag database restricted to the predicted true-positive taxa. Specifically, for the retained taxon set, VIP2B re-evaluates tag uniqueness by comparing theoretically digestible tags only within the set, rather than against the full reference collection. This reduction in search space substantially increases the number of unique 2B tags available per taxon, improving the stability of abundance estimation in the second-round mapping. Second-round mapping and relative abundance estimation VIP2B re-maps sample tags to the sample-specific unique-tag database and estimates taxonomic relative abundance by normalizing tag counts with respect to theoretical digestibility. For each detected taxon 𝑖 within the predicted true-positive taxa, its relative abundance is computed as 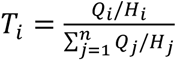.

### UHGV annotation-linked abundance summarization and multi-layer outputs

To extend taxonomic virus abundance profiles into interpretable, multi-layer virome readouts, we implemented an annotation-linked summarization step that maps each detected viral unit to UHGV^25^ metadata and aggregates abundances into biologically meaningful categories. Briefly, we join the virus-level abundance profile to a curated UHGV annotation resource that provides three complementary layers of information. First, UHGV metadata includes categorical phenotype descriptors, such as predicted lifestyle (e.g., virulent/lytic versus temperate/lysogenic), jumbo-phage status, and independent verification labels. Using these annotations, we construct phenotype-resolved virome profiles by summing the abundances of viruses sharing the same phenotype label within each sample, yielding compact summaries that quantify the relative representation of phenotypic classes. Second, UHGV provides hierarchical viral taxonomy assignments as well as predicted host taxonomies. We leverage these hierarchies to generate taxonomy-stratified abundance summaries for both viruses and their putative hosts by aggregating virus abundances at successive taxonomic ranks (from higher to lower ranks). This representation supports analyses at flexible resolution and facilitates cross-study comparisons where rank-level robustness is preferred. Third, UHGV links viral units to functional annotation terms, including UniRef90 protein families and higher-level functional clusters. We translate virus abundances into function-resolved profiles by assigning each virus’s abundance to all functional terms annotated to it and then summing contributions across viruses within each sample. For viruses annotated with multiple terms, the abundance contribution is propagated to each term. Functional profiles are subsequently normalized within each sample to ensure comparability across samples.

### Simulation data generation and taxonomic abundance benchmarking

To benchmark viral taxonomic profiling under controlled conditions, we generated simulated metagenomic datasets based on the UHGV catalog and compared VIP2B with two representative reference-based profilers, Sylph and Phanta. To ensure a fair comparison, the reference databases for all three methods were constructed from UHGV, thereby minimizing differences in theoretical detection capacity due to database composition. The Sylph database was built using “sylph sketch -c 100”, following the recommendation for small genomes^34^, and the Phanta database was prepared according to its official documentation for custom database construction^24^.

For the standard simulation, we first randomly selected viral species from UHGV, with species richness set to 200, 400, or 600 species per sample. For each selected species, a couple of genomes were randomly chosen. Genome-level relative abundances were then assigned using a log-normal distribution and normalized to sum to one per sample. These genome-level abundances were used as the ground truth for read simulation. Simulated metagenomic reads were generated from the selected genome FASTA sequences using wgsim (https://github.com/lh3/wgsim)^55^ to produce 150-bp paired-end reads, with a total sequencing depth of 10 million reads per sample. Genome-level abundances were subsequently summed to obtain the species-level ground truth abundance profile used for benchmarking against the species-level outputs from VIP2B, Sylph, and Phanta. For the dark-matter simulation, we followed the same overall procedure as in the standard simulation, except that only 80% of UHGV genomes were used to build the reference databases for all methods, while simulated reads were generated from genomes randomly selected from the remaining 20% that were excluded from database construction. The number of species was fixed at 200 per sample. For the incomplete-genome simulation, we again followed the same procedure as in the standard simulation, except that the genomes used for read simulation were truncated to 70%, 80%, or 90% of their original lengths before simulation. Species number was also fixed at 200 per sample. For the host-contamination simulation, viral reads were mixed with either bacterial or human reads at different host contamination ratios. For these datasets, the total sequencing depth was fixed at 12 million reads per sample, including both viral and host-derived reads. Because viral sequences can be embedded within bacterial genomes, we used geNomad v1.11.1^56^ to identify and remove prophage and other viral fragments from bacterial genomes before mixing bacterial reads with viral reads for benchmarking. This step ensured that the viral ground-truth profiles reflected only the abundances of the intended viral genomes in the simulated samples.

Each simulation condition included 10 independent replicates. Profiling performance was evaluated by comparing inferred species-level abundance profiles with the species-level ground truth using area under the precision-recall curve (AUPRC), F1 score, precision, and recall for taxon detection accuracy, and using Bray-Curtis similarity and L2 similarity for abundance concordance.

### Functional abundance benchmarking

To evaluate the accuracy of taxonomy-guided functional inference, we benchmarked VIP2B-inferred viral functional profiles using the standard simulation datasets described above. For each simulated sample, VIP2B first generated a taxonomic abundance profile, which was then converted into a UniRef90 gene family abundance profile through a genome content network (GCN). In this network, viral taxa were linked to their annotated gene families based on genome content information derived from UHGV, allowing gene family abundances to be inferred from the abundances of the corresponding viral taxa. In parallel, for each simulated sample, we constructed a custom viral UniRef90 protein database by collecting viral proteins from the viral taxa present in the ground-truth community. Simulated reads were then aligned against this database using DIAMOND v2.1.10^57^ with the options “--fast --iterate -k 1 --evalue 1e-5”, and the resulting mappings were used to derive the observed UniRef90 gene family abundance profile. This direct-mapping-derived profile was used as the reference functional profile for benchmarking. The inferred gene family profile generated from VIP2B taxonomic abundances was then compared with the observed gene family profile obtained by direct mapping from the same simulated sample.

### Metagenomic sample collection and processing

We assembled three metagenomic data collections for this study: one for paired bulk and virus-like particle (VLP)-enriched metagenomic sample comparisons, one for diversity, differential, and predictive analyses, and one for large-scale gut virome community type analysis. Dataset details are provided in Supplementary Table 2.

For the bulk-VLP comparison, we collected six studies with paired bulk and VLP metagenomes, including two infant cohorts and four adult cohorts. The infant datasets comprised PRJNA715601 and PRJEB46943 from the Copenhagen Prospective Studies on Asthma in Childhood (COPSAC; N=1,292)^58,59^, and PRJNA524703 from an early-life gut study of delivery mode (N=276)^60^. The adult datasets included PRJNA835720 from the Chinese Gut Virome Catalog (CHGV; N=326)^61^, PRJNA648796 and PRJNA648797 from an obesity study (N=456)^62^, PRJNA755142 from a hepatocellular carcinoma (HCC) study (N=24)^63^, and PRJEB57953 together with PRJEB57952 from a myalgic encephalomyelitis/chronic fatigue syndrome (ME/CFS) study (N=24)^64^.

For alpha-diversity, beta-diversity, differential, and machine learning analyses, we assembled 3,685 samples from 20 datasets spanning 16 disease or dysbiosis contexts, including 12 bulk-only studies and 4 paired bulk-VLP studies. The paired datasets were the delivery mode, obesity, HCC, and ME/CFS studies described above. The bulk-only datasets comprised ulcerative colitis and Crohn’s disease from PRJNA400072 (73 UC, 84 CD, 56 controls)^40^, colorectal cancer from PRJNA763023 (97 cases, 97 controls)^65^, multiple sclerosis from PRJEB28543 (24 cases, 24 controls)^66^, urinary tract infection from PRJNA400628 (312 cases, 220 controls)^67^, liver cirrhosis from PRJEB6337 (152 cases, 135 controls)^68^, atherosclerotic cardiovascular disease from PRJEB21528 (214 cases, 108 controls)^69^, type 2 diabetes from PRJNA422434 (186 cases, 183 controls)^70^, rheumatoid arthritis from PRJEB6997 (102 cases, 61 controls)^71^, Parkinson’s disease from PRJEB53401 (48 cases, 70 controls)^72^, and constipation- and diarrhea-predominant irritable bowel syndrome from PRJEB37924 (136 IBS-C, 169 IBS-D, 149 controls)^73^. For machine learning analyses, we included only datasets with more than 25 samples in both case and control groups; thus, all datasets were retained except HCC, ME/CFS, and multiple sclerosis.

For the gut virome community type analysis, we compiled 6,090 samples by combining the above-mentioned datasets, excluding the COPSAC and delivery mode datasets because they were derived from infants, with the CHGV dataset and stool samples from five additional cohorts: PRJDB4176 (N=286)^74^, PRJEB11532 (N=900)^75^, PRJEB39223 (N=568)^76^, PRJNA319574 (N=385)^77^, and PRJNA48479 (N=425)^78^.

Host-derived reads were not removed before profiling. VIP2B and MAP2B were designed to tolerate substantial human DNA background, enabling direct analysis of raw metagenomic datasets and substantially accelerating large-scale processing.

### Compare paired bulk and VLP metagenomic samples

Shared species were defined as those with non-zero relative abundance in both bulk and VLP profiles within the same matched sample. The abundances of these overlapping species were then pooled across matched samples in each dataset, log10-transformed, and compared between bulk and VLP data using a two-sample Kolmogorov-Smirnov test. P values were adjusted across datasets using the Benjamini-Hochberg method.

### Alpha and beta diversity analysis

Shannon and Simpson indices were used to quantify alpha diversity. Correlations between bacteriome and virome alpha diversity were assessed using Pearson correlation coefficients. For beta-diversity analysis, Bray–Curtis distance matrices were calculated from the abundance profiles, and permutational multivariate analysis of variance (PERMANOVA) was performed using the adonis2 function in the R package vegan to obtain P values, F statistics, and R^2^ values.

### Differential abundance and phenotype-associated viral feature analysis

Differential analyses of case-control differences were performed using MaAsLin3^79^ with disease status as the fixed effect and the healthy group as the reference level. For abundance-based taxonomic and functional profiles, input tables were total-sum scaled and log transformed before model fitting. For taxonomic coverage profiles, no additional normalization or transformation was applied.

To examine phenotype-associated viral features, we analyzed VIP2B-derived virulent phage abundance and jumbo phage abundance across disease cohorts. Group differences for each feature within each disease panel were assessed using two-sided Wilcoxon rank-sum tests.

### Machine learning predictive models

We constructed supervised case–control prediction models using multiple microbial feature sets, including VIP2B taxonomic, coverage, and functional profiles, Sylph species profiles, MAP2B taxonomic profiles, and combined feature representations. For each feature table, data were converted to sample-level matrices and features present in at least 10% of samples were retained. Feature names were labeled by data type to avoid naming conflicts after merging, and all models were trained and evaluated on the same intersected sample set. To avoid information leakage, all preprocessing steps were performed within each training fold only. Zero-variance features were removed using the training data, and feature selection was performed with Boruta (maxRuns = 50) with tentative features resolved by “TentativeRoughFix”^80^. The selected features were then used to train both Elastic Net and Random Forest models. Elastic Net models were fitted using “cv.glmnet” with a binomial family and 5-fold internal cross-validation^81^, whereas Random Forest models were trained using the “randomForest” R package^82^ with training-set-based tuning of “mtry”. For the combined-feature model, features selected from the individual base feature sets were pooled within each training fold and used directly for model fitting.

Model performance was evaluated using 5 repeats of stratified 10-fold cross-validation, with identical train-test splits applied across feature sets within each repeat to ensure fair comparison. In each fold, models were trained on nine folds and evaluated on the held-out fold. Predictive performance was summarized using the area under the receiver operating characteristic curve (AUROC), reported as the mean, standard deviation, and 95% confidence interval across repeats. ROC curves were generated from held-out predictions.

### Gut virome community type identification

Input matrices were formatted as sample-by-species tables, with duplicate sample IDs removed, missing values set to zero, all-zero species discarded, and samples with no remaining signal excluded. For distance-based clustering, profiles were converted to relative abundances after filtering low-prevalence features, using a 1% prevalence threshold for taxonomic and coverage profiles and a 10% threshold for functional profiles. Partitioning around medoids (PAM) clustering was then performed for K=2 to 10 using Bray-Curtis, Jensen-Shannon, and root Jensen-Shannon distances. Cluster validity was assessed using the Calinski-Harabasz index, prediction strength, and mean silhouette score, with higher values indicating stronger support for a given number of clusters. As a complementary model-based approach, Dirichlet multinomial mixture (DMM) models were fitted to coverage, abundance, and functional profiles for K=1 to 10, and model fit was evaluated using the Bayesian Information Criterion (BIC), with lower values indicating better support for the corresponding number of clusters.

After virome community types were defined, viral alpha diversity within each community type was quantified using the Shannon index and compared using two-sided Wilcoxon rank-sum tests. Differences in the proportions of healthy and disease samples between community types were assessed using two-sided Fisher’s exact tests.

### Data availability

Raw metagenomic reads were downloaded from the Sequence Read Archive (SRA). The corresponding accession numbers and metadata are provided in Supplementary Table 2. The bulk and VLP sample-pairing information for the COPSAC dataset has been removed in accordance with strict GDPR regulations regarding the publication of individual-level data in Europe.

### Code availability

VIP2B software and its complete instructions for installation and usage are available at https://github.com/sunzhengCDNM/VIP2B/tree/perf/accelerated-version.

## Acknowledgments

We gratefully thank the COPSAC team, especially Dr. Shiraz Shah and Dr. Simon Rasmussen, for providing the bulk and VLP sample-pairing information for the COPSAC dataset. We also thank Rong Xia and Dr. Peipei Gao for their assistance with software testing.

## REFERENCES

1. Hendrix, R. W., Smith, M. C. M., Burns, R. N., Ford, M. E. & Hatfull, G. F. Evolutionary relationships among diverse bacteriophages and prophages: All the world’s a phage. Proc. Natl. Acad. Sci. U.S.A. 96, 2192–2197 (1999).

2. Stern, A. & Sorek, R. The phage-host arms race: Shaping the evolution of microbes. BioEssays 33, 43–51 (2011).

3. Cesar Ignacio-Espinoza, J., Solonenko, S. A. & Sullivan, M. B. The global virome: not as big as we thought? Current Opinion in Virology 3, 566–571 (2013).

4. Kurilovich, E. & Geva-Zatorsky, N. Effects of bacteriophages on gut microbiome functionality. Gut Microbes 17, 2481178 (2025).

5. Rodriguez-Valera, F. et al. Explaining microbial population genomics through phage predation. Nat Rev Microbiol 7, 828–836 (2009).

6. Touchon, M., Moura De Sousa, J. A. & Rocha, E. P. Embracing the enemy: the diversification of microbial gene repertoires by phage-mediated horizontal gene transfer. Current Opinion in Microbiology 38, 66–73 (2017).

7. Mahmud, Md. R., et al. Role of bacteriophages in shaping gut microbial community. Gut Microbes 16, 2390720 (2024).

8. Gonzalez Pastor, B., Shkoporov, A. N. & Hill, C. Not just passengers: Phages as agents of genetic exchange in fecal microbiota transplantation. Cell Host & Microbe S1931312826001253 (2026) doi:10.1016/j.chom.2026.03.017.

9. Norman, J. M. et al. Disease-Specific Alterations in the Enteric Virome in Inflammatory Bowel Disease. Cell 160, 447–460 (2015).

10. Gogokhia, L. et al. Expansion of Bacteriophages Is Linked to Aggravated Intestinal Inflammation and Colitis. Cell Host & Microbe 25, 285–299.e8 (2019).

11. Lloyd-Price, J. et al. Multi-omics of the gut microbial ecosystem in inflammatory bowel diseases. Nature 569, 655–662 (2019).

12. Majzoub, M. E. et al. The phageome of patients with ulcerative colitis treated with donor fecal microbiota reveals markers associated with disease remission. Nat Commun 15, 8979 (2024).

13. Wen, W. et al. Gut DNA virome enterotype dictates inflammation heterogeneity through tuning the phage-bacteria-sphingosine-intestine axis in Crohn’s disease. Cell Host & Microbe 34, 460–478.e9 (2026).

14. De Jonge, P. A. et al. Gut virome profiling identifies a widespread bacteriophage family associated with metabolic syndrome. Nat Commun 13, 3594 (2022).

15. Cervantes-Echeverría, M., Gallardo-Becerra, L., Cornejo-Granados, F. & Ochoa-Leyva, A. The Two-Faced Role of crAssphage Subfamilies in Obesity and Metabolic Syndrome: Between Good and Evil. Genes 14, 139 (2023).

16. Mao, X. et al. Transfer of modified gut viromes improves symptoms associated with metabolic syndrome in obese male mice. Nat Commun 15, 4704 (2024).

17. Deng, Y., Jiang, S., Duan, H., Shao, H. & Duan, Y. Bacteriophages and their potential for treatment of metabolic diseases. Journal of Diabetes 16, e70024 (2024).

18. Reyes, A., Semenkovich, N. P., Whiteson, K., Rohwer, F. & Gordon, J. I. Going viral: next-generation sequencing applied to phage populations in the human gut. Nat Rev Microbiol 10, 607–617 (2012).

19. Roux, S. et al. Minimum Information about an Uncultivated Virus Genome (MIUViG). Nat Biotechnol 37, 29–37 (2019).

20. Camarillo-Guerrero, L. F., Almeida, A., Rangel-Pineros, G., Finn, R. D. & Lawley, T. D. Massive expansion of human gut bacteriophage diversity. Cell 184, 1098–1109.e9 (2021).

21. Sutton, T. D. S., Clooney, A. G., Ryan, F. J., Ross, R. P. & Hill, C. Choice of assembly software has a critical impact on virome characterisation. Microbiome 7, 12 (2019).

22. Wang, H. et al. Complementary insights into gut viral genomes: a comparative benchmark of short- and long-read metagenomes using diverse assemblers and binners. Microbiome 12, 260 (2024).

23. Lu, J. et al. Metagenome analysis using the Kraken software suite. Nat Protoc 17, 2815–2839 (2022).

24. Pinto, Y., Chakraborty, M., Jain, N. & Bhatt, A. S. Phage-inclusive profiling of human gut microbiomes with Phanta. Nat Biotechnol 42, 651–662 (2024).

25. Camargo, A. P. et al. A genomic atlas of the human gut virome elucidates genetic factors shaping host interactions. Preprint at 10.1101/2025.11.01.686033 (2025).

26. Nayfach, S. et al. CheckV assesses the quality and completeness of metagenome-assembled viral genomes. Nat Biotechnol 39, 578–585 (2021).

27. Jurasz, H., Pawłowski, T. & Perlejewski, K. Contamination Issue in Viral Metagenomics: Problems, Solutions, and Clinical Perspectives. Front. Microbiol. 12, 745076 (2021).

28. Kim, N. et al. Genome-resolved metagenomics: a game changer for microbiome medicine. Exp Mol Med 56, 1501–1512 (2024).

29. Tegegne, H. A. & Savidge, T. C. Gut microbiome metagenomics in clinical practice: bridging the gap between research and precision medicine. Gut Microbes 17, 2569739 (2025).

30. Sun, Z. et al. Species-resolved sequencing of low-biomass or degraded microbiomes using 2bRAD-M. Genome Biol 23, 36 (2022).

31. Sun, Z. et al. Removal of false positives in metagenomics-based taxonomy profiling via targeting Type IIB restriction sites. Nat Commun 14, 5321 (2023).

32. Marshall, J. J. T. & Halford, S. E. The Type IIB restriction endonucleases. Biochemical Society Transactions 38, 410–416 (2010).

33. Wang, S., Meyer, E., McKay, J. K. & Matz, M. V. 2b-RAD: a simple and flexible method for genome-wide genotyping. Nat Methods 9, 808–810 (2012).

34. Shaw, J. & Yu, Y. W. Rapid species-level metagenome profiling and containment estimation with sylph. Nat Biotechnol 10.1038/s41587-024-02412-y (2024) doi:10.1038/s41587-024-02412-y.

35. Gregory, A. C. et al. The Gut Virome Database Reveals Age-Dependent Patterns of Virome Diversity in the Human Gut. Cell Host & Microbe 28, 724–740.e8 (2020).

36. Johansen, J. et al. Genome binning of viral entities from bulk metagenomics data. Nat Commun 13, 965 (2022).

37. Li, Y. et al. Biases and complementarity in gut viromes obtained from bulk and virus-like particle-enriched metagenomic sequencing. Microbiol Spectr e00013–25 (2025) doi:10.1128/spectrum.00013-25.

38. Maqsood, R. et al. Discordant transmission of bacteria and viruses from mothers to babies at birth. Microbiome 7, 156 (2019).

39. Zeng, S. et al. A metagenomic catalog of the early-life human gut virome. Nat Commun 15, 1864 (2024).

40. Franzosa, E. A. et al. Gut microbiome structure and metabolic activity in inflammatory bowel disease. Nat Microbiol 4, 293–305 (2018).

41. Clooney, A. G. et al. Whole-Virome Analysis Sheds Light on Viral Dark Matter in Inflammatory Bowel Disease. Cell Host & Microbe 26, 764–778.e5 (2019).

42. Park, H. et al. Cross-cohort analysis reveals conserved gut virome signatures and phage-encoded auxiliary functions in ulcerative colitis. Preprint at 10.64898/2026.01.21.700755 (2026).

43. Jensen, J. S. L. et al. Enhanced multi-omic viral profiling from microbial community sequencing with BAQLaVa. Preprint at 10.64898/2026.02.11.705346 (2026).

44. Zhang, S. et al. Prophages and their interactions with lytic phages in the human gut microbiota and their impact on microbial diversity, gut health, and disease. Appl Environ Microbiol 91, e01899–25 (2025).

45. Chen, L. et al. Animal-associated jumbo phages as widespread and active modulators of gut microbiome ecology and metabolism. Sci. Adv. 12, eaeb6265 (2026).

46. Wheatley, R. M., Holtappels, D. & Koskella, B. Evaluation of bacteriophages as a signature of microbiome health: a systematic review and meta-analysis. The Lancet Microbe 6, 101196 (2025).

47. Caliński, T. & Harabasz, J. A dendrite method for cluster analysis. Communications in Statistics 3, 1–27 (1974).

48. Tibshirani, R. & Walther, G. Cluster Validation by Prediction Strength. Journal of Computational and Graphical Statistics 14, 511–528 (2005).

49. Rousseeuw, P. J. Silhouettes: A graphical aid to the interpretation and validation of cluster analysis. Journal of Computational and Applied Mathematics 20, 53–65 (1987).

50. Koren, O. et al. A Guide to Enterotypes across the Human Body: Meta-Analysis of Microbial Community Structures in Human Microbiome Datasets. PLoS Comput Biol 9, e1002863 (2013).

51. Holmes, I., Harris, K. & Quince, C. Dirichlet Multinomial Mixtures: Generative Models for Microbial Metagenomics. PLoS ONE 7, e30126 (2012).

52. Song, L., Zhang, L. & Fang, X. Characterizing Enterotypes in Human Metagenomics: A Viral Perspective. Front. Microbiol. 12, 740990 (2021).

53. Jansen, D. et al. Community Types of the Human Gut Virome are Associated with Endoscopic Outcome in Ulcerative Colitis. Journal of Crohn’s and Colitis 17, 1504–1513 (2023).

54. Qingbo, L. et al. Identification of enterotype and its predictive value for patients with colorectal cancer. Gut Pathog 16, 12 (2024).

55. Li, H. et al. The Sequence Alignment/Map format and SAMtools. Bioinformatics 25, 2078–2079 (2009).

56. Camargo, A. P. et al. Identification of mobile genetic elements with geNomad. Nat Biotechnol 42, 1303–1312 (2024).

57. Buchfink, B., Xie, C. & Huson, D. H. Fast and sensitive protein alignment using DIAMOND. Nat Methods 12, 59–60 (2015).

58. Shah, S. A. et al. Expanding known viral diversity in the healthy infant gut. Nat Microbiol 8, 986–998 (2023).

59. Li, X. et al. The infant gut resistome associates with E. coli, environmental exposures, gut microbiome maturity, and asthma-associated bacterial composition. Cell Host & Microbe 29, 975–987.e4 (2021).

60. Liang, G. et al. The stepwise assembly of the neonatal virome is modulated by breastfeeding. Nature 581, 470–474 (2020).

61. Chen, J. et al. Efficient Recovery of Complete Gut Viral Genomes by Combined Short- and Long-Read Sequencing. Advanced Science 11, 2305818 (2024).

62. Yang, K. et al. Alterations in the Gut Virome in Obesity and Type 2 Diabetes Mellitus. Gastroenterology 161, 1257–1269.e13 (2021).

63. Zhang, F. et al. Critical Assessment of Whole Genome and Viral Enrichment Shotgun Metagenome on the Characterization of Stool Total Virome in Hepatocellular Carcinoma Patients. Viruses 15, 53 (2022).

64. Hsieh, S.-Y. et al. Investigating the Human Intestinal DNA Virome and Predicting Disease-Associated Virus–Host Interactions in Severe Myalgic Encephalomyelitis/Chronic Fatigue Syndrome (ME/CFS). IJMS 24, 17267 (2023).

65. Yang, Y. et al. Dysbiosis of human gut microbiome in young-onset colorectal cancer. Nat Commun 12, 6757 (2021).

66. Ventura, R. E. et al. Gut microbiome of treatment-naïve MS patients of different ethnicities early in disease course. Sci Rep 9, 16396 (2019).

67. Worby, C. J. et al. Longitudinal multi-omics analyses link gut microbiome dysbiosis with recurrent urinary tract infections in women. Nat Microbiol 7, 630–639 (2022).

68. Qin, N. et al. Alterations of the human gut microbiome in liver cirrhosis. Nature 513, 59–64 (2014).

69. Jie, Z. et al. The gut microbiome in atherosclerotic cardiovascular disease. Nat Commun 8, 845 (2017).

70. Qin, J. et al. A metagenome-wide association study of gut microbiota in type 2 diabetes. Nature 490, 55–60 (2012).

71. Guo, R. et al. Dysbiotic Oral and Gut Viromes in Untreated and Treated Rheumatoid Arthritis Patients. Microbiol Spectr 10, e00348–22 (2022).

72. Boktor, J. C. et al. Integrated Multi-Cohort Analysis of the Parkinson’s Disease Gut Metagenome. Movement Disorders 38, 399–409 (2023).

73. Mars, R. A. T. et al. Longitudinal Multi-omics Reveals Subset-Specific Mechanisms Underlying Irritable Bowel Syndrome. Cell 182, 1460–1473.e17 (2020).

74. Yachida, S. et al. Metagenomic and metabolomic analyses reveal distinct stage-specific phenotypes of the gut microbiota in colorectal cancer. Nat Med 25, 968–976 (2019).

75. Zeevi, D. et al. Personalized Nutrition by Prediction of Glycemic Responses. Cell 163, 1079–1094 (2015).

76. Asnicar, F. et al. Microbiome connections with host metabolism and habitual diet from 1,098 deeply phenotyped individuals. Nat Med 27, 321–332 (2021).

77. Schirmer, M. et al. Linking the Human Gut Microbiome to Inflammatory Cytokine Production Capacity. Cell 167, 1125–1136.e8 (2016).

78. Lloyd-Price, J. et al. Strains, functions and dynamics in the expanded Human Microbiome Project. Nature 550, 61–66 (2017).

79. Nickols, W. A. et al. MaAsLin 3: refining and extending generalized multivariable linear models for meta-omic association discovery. Nat Methods 23, 554–564 (2026).

80. Kursa, M. B. & Rudnicki, W. R. Feature Selection with the Boruta Package. J. Stat. Soft. 36, (2010).

81. Tay, J. K., Narasimhan, B. & Hastie, T. Elastic Net Regularization Paths for All Generalized Linear Models. J. Stat. Soft. 106, (2023).

82. Breiman, L. Random Forests. Machine Learning 45, 5–32 (2001).

