## Supplementary Figures 1-8 for "The human gut virome is a non-redundant and clinically informative component of the microbiome"

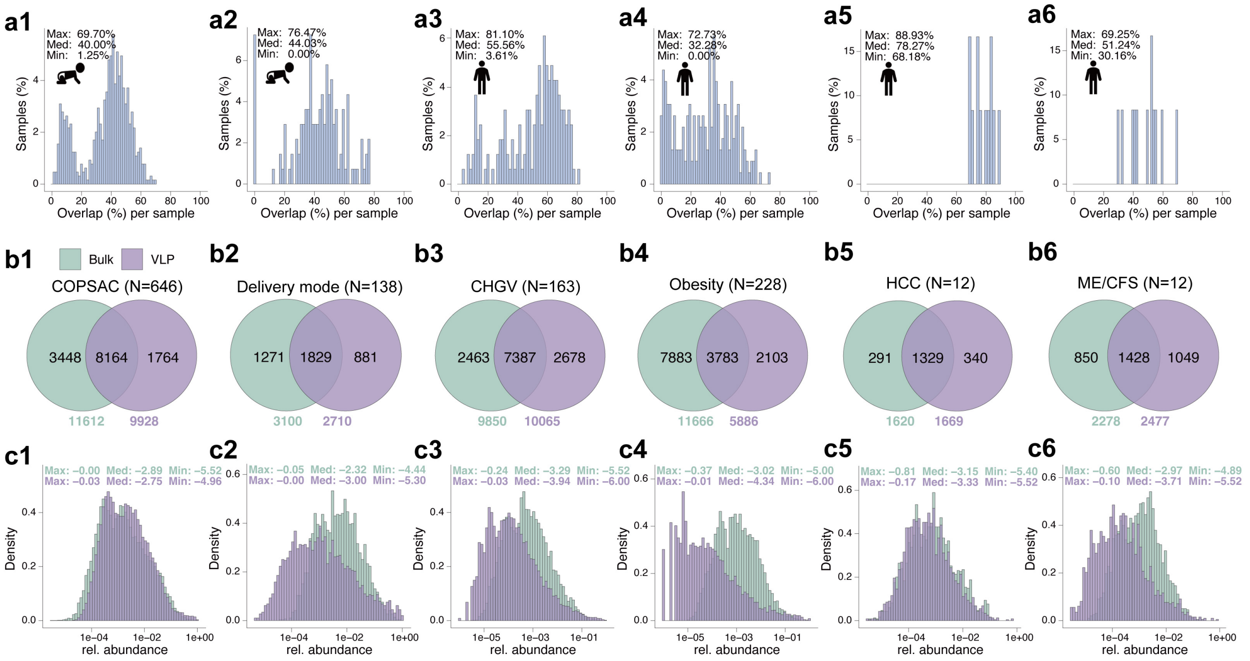


**Figure S1.** Overlap of viral species between paired bulk shotgun and VLP-enriched metagenomes using Sylph across six cohorts. Number 1-6 indicate two infant cohorts: COPSAC (Copenhagen Prospective Studies on Asthma in Childhood) and delivery mode, and four adult cohorts: CHGV (Chinese Gut Virome Catalog), obesity, HCC, (hepatocellular carcinoma) and ME/CFS (myalgic encephalomyelitis/chronic fatigue syndrome). (**a1-a6**) Distributions of the percentage of VLP-detected viral species that are also detected in the paired bulk metagenome for each sample. Minimum (Min), median (Med), and maximum (Max) of the percentages are annotated. (**b1-b6**) Venn diagrams showing the numbers of shared and method-specific viral species recovered from bulk and VLP data within each cohort. (**c1-c6**) Density distributions of cumulative abundances of overlapping viral species in bulk and VLP data. Minimum (Min), median (Med), and maximum (Max) of the log10-scaled abundances are annotated.

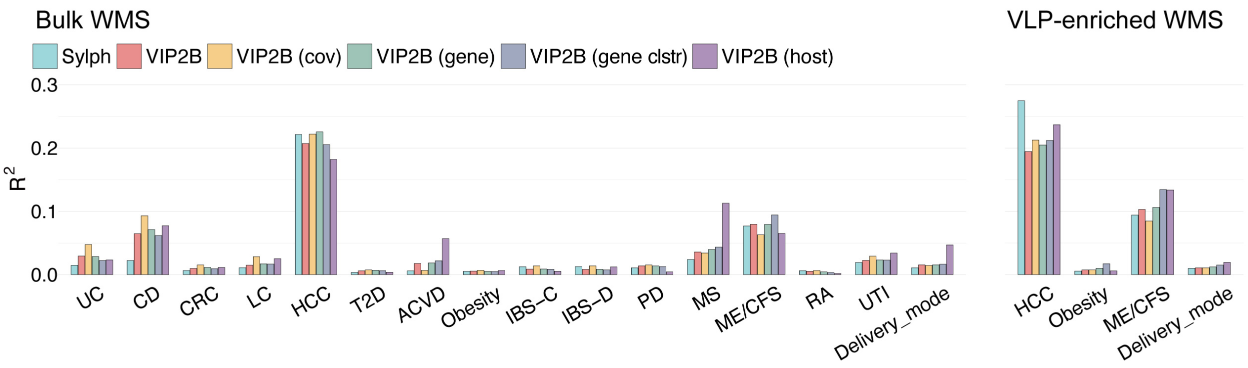


**Figure S2.** $R^2$ values of the permutational multivariate analysis of variance (PERMANOVA, Adonis permutations=9999) comparing beta diversity between controls and dysbiosis using VIP2B-derived profiles.

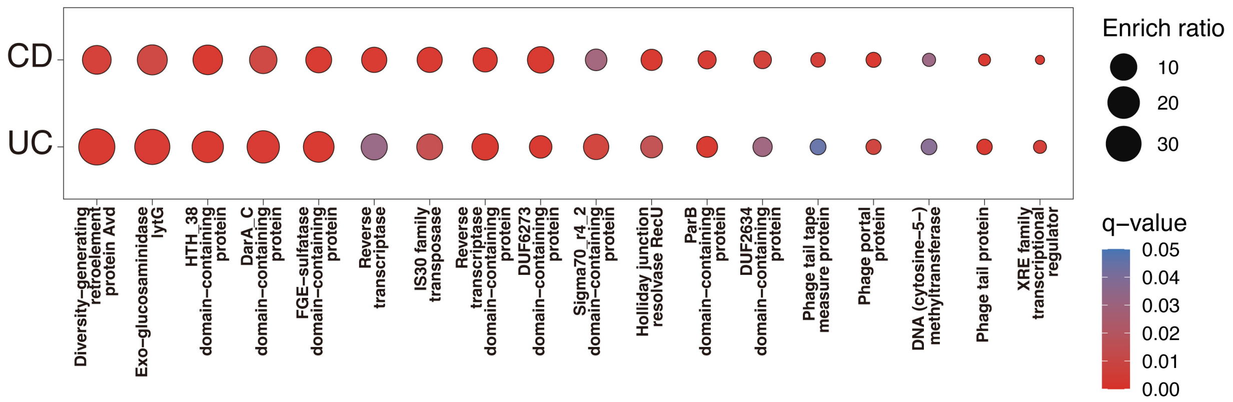


**Figure S3.** Viral gene clusters consistently depleted in both Crohn's disease (CD) and ulcerative colitis (UC) are enriched for functions related to bacterial host recognition and prophage maintenance.

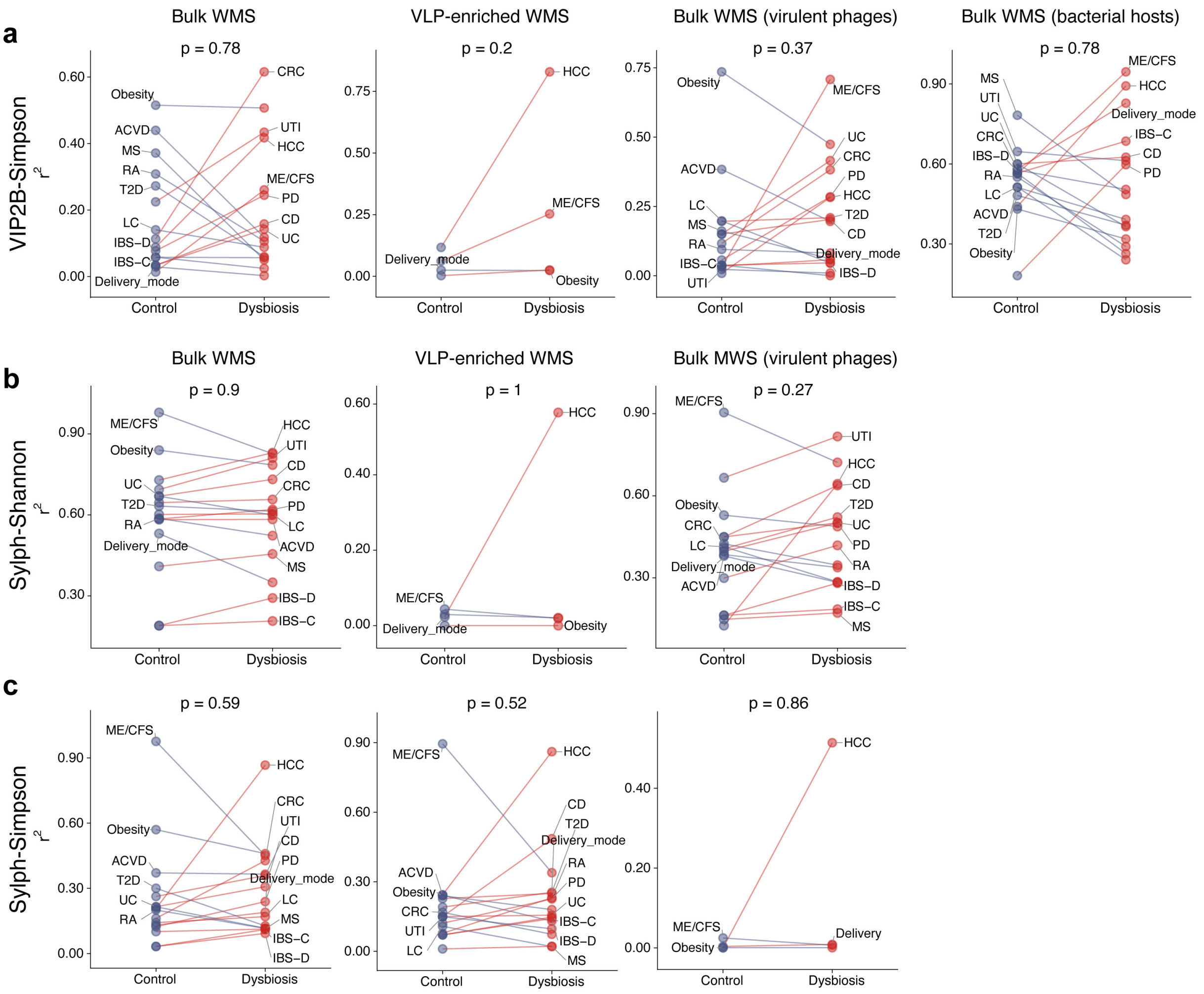


**Figure S4.** Bacteriome-virome alpha-diversity correlations are not consistently reduced in dysbiosis across datasets and sequencing types using Simpson index. (**a-b**) Pearson correlation coefficients of determination ($r^2$) between bacterial and viral Simpson indices in controls versus dysbiosis, computed using species-level viral taxonomic abundance profiles generated by (**a**) VIP2B and (**b**) Sylph. Results are shown for 16 bulk metagenomic datasets (left), 4 VLP-enriched metagenomic datasets (middle left), 16 bulk metagenomic datasets with viruses limited to lytic phages (middle right), and 16 bulk metagenomic datasets with bacteria limited to the hosts of phages (right). When the correlation is higher in controls than in dysbiosis, the dysbiosis condition is labeled on the left and the connecting line is blue. When the correlation is higher in dysbiosis than in controls, the dysbiosis condition is labeled on the right and the connecting line is red.

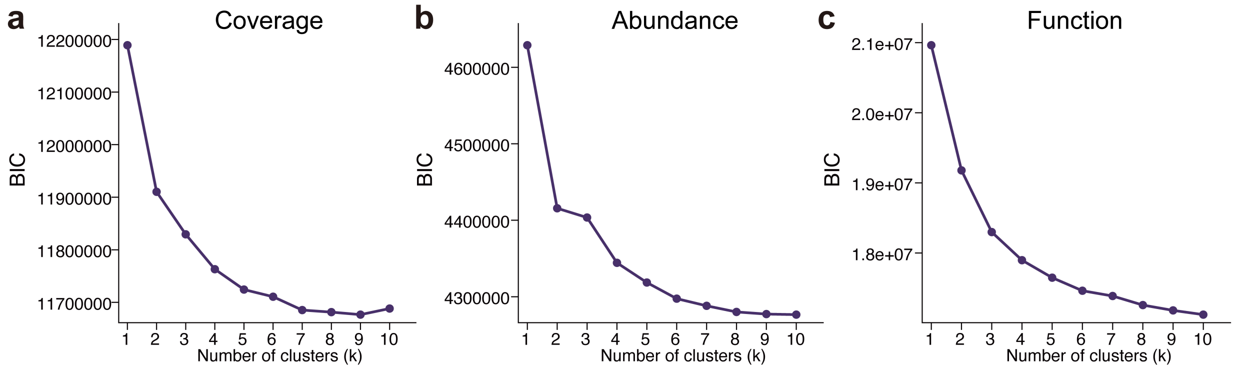


**Figure S5.** Cluster-number selection for virome profiles. (**a-c**) Bayesian Information Criterion (BIC) (lower is better) from Dirichlet multinomial mixture (DMM) models for (**a**) coverage, (**b**) abundance and (**c**) functional profiles across K = 1-10.

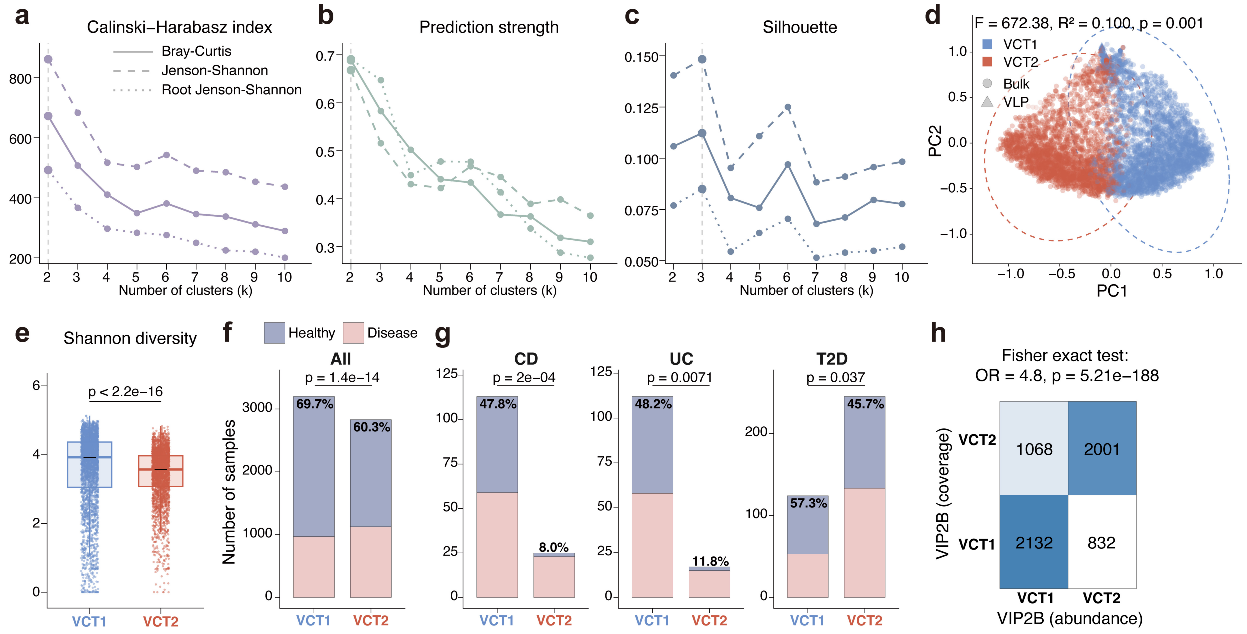


**Figure S6**. VIP2B abundance profiles from 6,090 stool metagenomic samples reveal two virome enterotypes. (**a-c**) Calinski-Harabasz (CH) index (**a**), prediction strength (**b**) and Silhouette score (**c**) for partitioning around medoids (PAM) clustering using Bray-Curtis, Jensen-Shannon, and root Jensen-Shannon distances on species-level taxonomic coverage profiles, evaluated across K = 2-10 clusters. (**d**) Principal coordinates analysis (PCoA) show separation of two virome community types (VCTs), VCT1 (blue) and VCT2 (red), supported by PERMANOVA $p < 0.05$ (Adonis, permutations = 999). Bulk and VLP metagenomic samples are shown as circles and triangles, respectively. (**e**) Viral alpha diversity (Shannon index) is higher in VCT1 than in VCT2 (p value was calculated using a two-sided Wilcoxon rank-sum test). (**f-g**) Percentages of healthy and disease samples in VCT1 and VCT2 (**f**) across all samples and (**g**) within disease cohorts. The stacked bar plots also show the percentage of healthy samples within each enterotype. P values were computed using two-sided Fisher's exact tests. Disease cohorts include Crohn's disease (CD), ulcerative colitis (UC), and type 2 diabetes (T2D). (**h**) Contingency analysis of enterotype assignments from coverage and abundance profiles. Significance of agreement was evaluated using Fisher's exact test. OR, odds ratio.

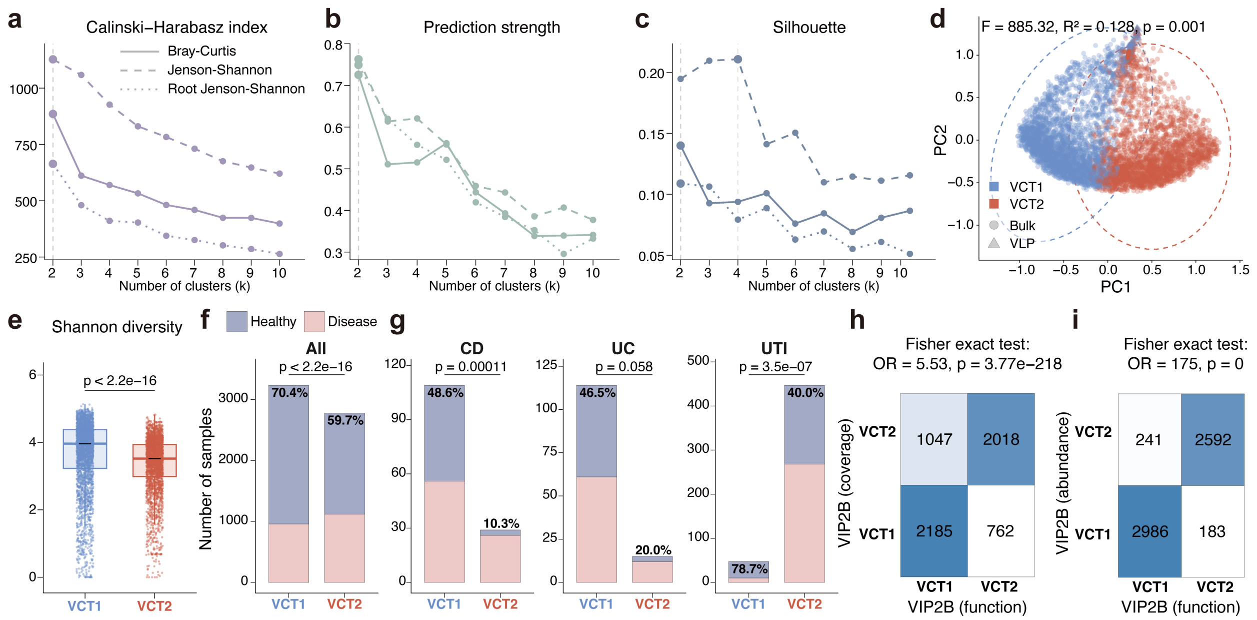


**Figure S7**. VIP2B functional (gene level) profiles from 6,090 stool metagenomic samples reveal two virome enterotypes. (**a-c**) Calinski-Harabasz (CH) index (**a**), prediction strength (**b**) and Silhouette score (**c**) for partitioning around medoids (PAM) clustering using Bray-Curtis, Jensen-Shannon, and root Jensen-Shannon distances on species-level taxonomic coverage profiles, evaluated across K = 2-10 clusters. (**d**) Principal coordinates analysis (PCoA) show separation of two virome community types (VCTs), VCT1 (blue) and VCT2 (red), supported by PERMANOVA $p < 0.05$ (Adonis, permutations = 999). Bulk and VLP metagenomic samples are shown as circles and triangles, respectively. (**e**) Viral alpha diversity (Shannon index) is higher in VCT1 than in VCT2 (p value was calculated using a two-sided Wilcoxon rank-sum test). (**f-g**) Percentages of healthy and disease samples in VCT1 and VCT2 (**f**) across all samples and (**g**) within Crohn's disease (CD), ulcerative colitis (UC) and urinary tract infection (UTI) cohorts. The stacked bar plots also show the percentage of healthy samples within each enterotype. P values were computed using two-sided Fisher's exact tests. (**h-i**) Contingency analysis of enterotype assignments from (**h**) coverage versus function profiles and (**i**) abundance versus function profiles. Significance of agreement was evaluated using Fisher's exact test. OR, odds ratio.

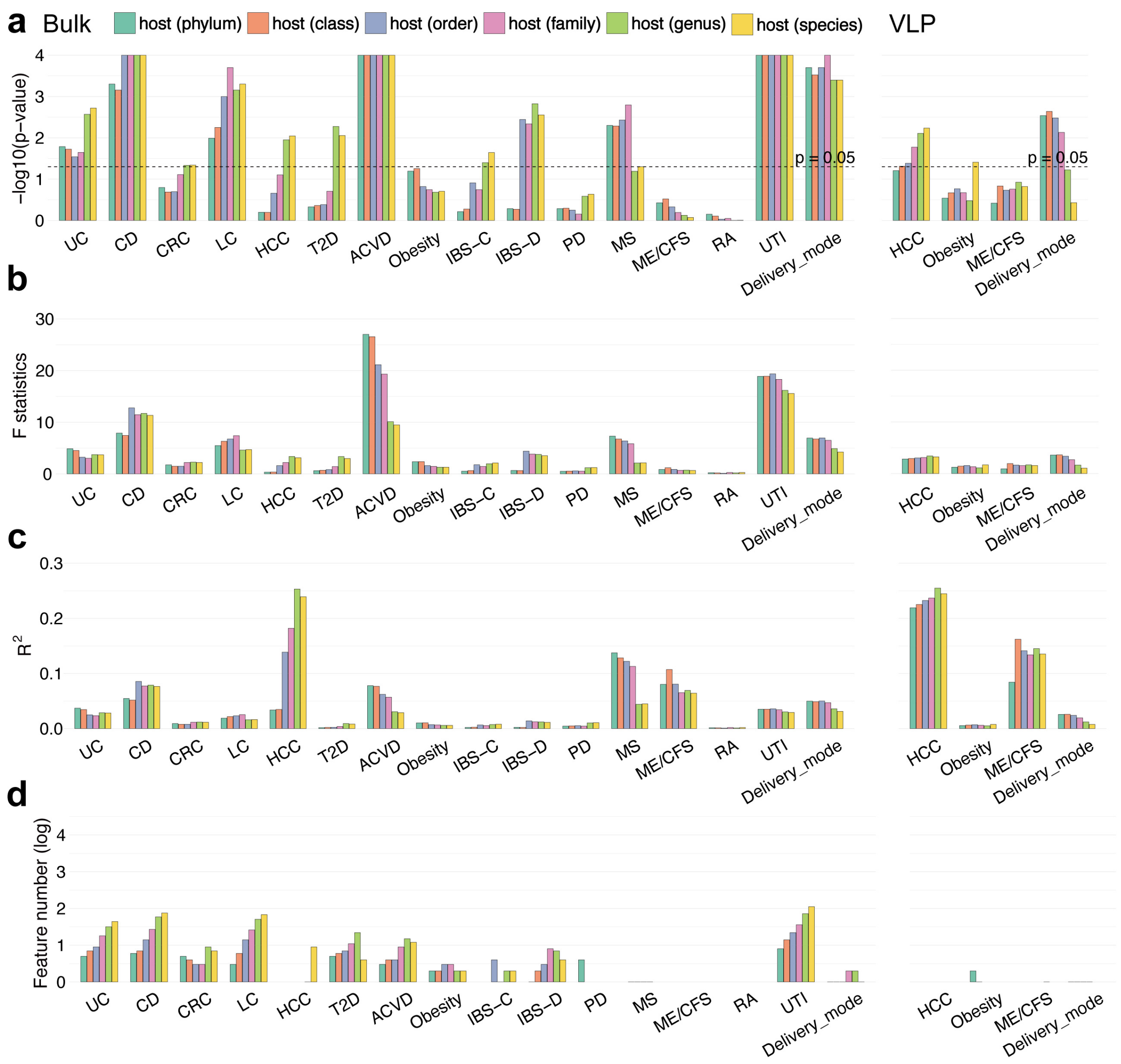


**Figure S8.** Viral host compositional differences and dysbiosis-associated viral signatures at different taxonomic levels across 20 datasets. (**a-c**) Permutational multivariate analysis of variance (PERMANOVA, ADONIS permutations=9999) comparing beta diversity between controls and dysbiosis using VIP2B-derived profiles, shown as (**a**) p values, (**b**) F statistics, and (**c**) $R^2$. Virus host taxonomic abundance profiles include abundances at phylum, class, order, family, genus and species levels. (**d**) Numbers of dysbiosis-associated viral features (FDR-adjusted $p < 0.05$) identified by MaAsLin3.
